# Relocation of rDNA repeats for repair is dependent on SUMO-mediated nucleolar release by the Cdc48/p97 segregase

**DOI:** 10.1101/2021.01.05.425376

**Authors:** Matías Capella, Imke K. Mandemaker, Fabian den Brave, Lucía Martín Caballero, Boris Pfander, Andreas G. Ladurner, Stefan Jentsch, Sigurd Braun

## Abstract

Ribosomal RNA genes (rDNA) are highly unstable and susceptible to rearrangement due to active transcription and their repetitive nature. Compartmentalization of rDNA in the nucleolus suppresses uncontrolled recombination. However, broken repeats must be released to the nucleoplasm to allow repair by homologous recombination. The process of rDNA relocation is conserved from yeast to humans, but the underlying molecular mechanisms are currently unknown. Here we show that DNA damage induces phosphorylation of the CLIP-cohibin complex, releasing membrane-tethered rDNA from the nucleolus in *Saccharomyces cerevisiae*. Downstream of phosphorylation, SUMOylation targets CLIP-cohibin for disassembly mediated by the Cdc48/p97 chaperone, which recognizes SUMOylated CLIP-cohibin through its cofactor, Ufd1. Consistent with a conserved mechanism, UFD1L depletion impairs rDNA release in human cells. The dynamic and regulated assembly and disassembly of the rDNA-tethering complex is therefore a key determinant of nucleolar rDNA release and genome integrity.

## Introduction

Eukaryotic genomes contain large amounts of repetitive sequences at centromeres, telomeres, and ribosomal RNA genes (rDNA). In *Saccharomyces cerevisiae*, the rDNA locus consists of approximately 150 copies, organized in 9.1 kb tandem repeats on chromosome XII. Humans have hundreds of 43 kb rDNA units located on different chromosomes. This large availability of donor sequences means that rDNA repeats often undergo homologous recombination (HR), making them a major source of genome instability (Kobayashi, 2011). As an example, DNA double-strand break (DSB) repair within a repetitive array can cause repeat insertions or deletions through HR with a neighboring unit (Jaco et al., 2008; Kobayashi, 2011). Such rearrangements can have severe consequences. Indeed, rDNA repeat translocations are associated with cellular senescence, neurodegenerative diseases, and are amongst the most common events observed in cancer cells (Hallgren et al., 2014; Lindström et al., 2018; Sinclair and Guarente, 1997; Stults et al., 2009).

The rDNA repeats are spatially segregated into the nucleolus, a membrane-less subnuclear organelle. In *S. cerevisiae*, rDNA is tethered to the nuclear envelope by the CLIP (<u>c</u>hromatin linkage of inner nuclear membrane <u>p</u>roteins) and the cohibin complexes (Mekhail et al., 2008). Cohibin is a “V”-shaped complex composed of one Lrs4 homodimer and two Csm1 homodimers. It associates with rDNA through its interaction with the RENT complex (<u>re</u>gulator of <u>n</u>ucleolar silencing and telophase exit) and Tof2 (Corbett et al., 2010; Huang et al., 2006; Mekhail et al., 2008). Cohibin tethers rDNA to the nuclear periphery by binding the CLIP complex. CLIP comprises two integral inner nuclear membrane proteins, the LEM domain protein Heh1 (also known as Src1) and Nur1 (Supplementary Figure 1a) (Mekhail et al., 2008).

SUMOylation plays multiple roles in DNA damage repair. Akin to ubiquitylation, an enzymatic cascade covalently attaches the small ubiquitin-like modifier SUMO to specific lysine residues on target proteins. SUMO and its conjugates are recognized by proteins containing SIMs (<u>S</u>UMO interacting <u>m</u>otifs) (Geiss-Friedlander and Melchior, 2007). Furthermore, several SUMO moieties can be conjugated to form a chain, which triggers protein degradation through the recruitment of SUMO-targeted ubiquitin ligases (STUbLs) (Prudden et al., 2007). Conversely, SUMO moieties can be removed by the SUMO-specific proteases Ulp1 and Ulp2, which also mediate SUMO maturation by exposing the C-terminal Gly-Gly motif (Geiss-Friedlander and Melchior, 2007).

SUMOylation of Rad52, a key player in DSB repair, is one of an array of mechanisms that suppresses aberrant rDNA repeat recombination (Mekhail et al., 2008; Torres-Rosell et al., 2007). Together with the Smc5/6 (<u>s</u>tructural <u>m</u>aintenance of <u>c</u>hromosomes) and MRX (Mre11-Rad50-Xrx2) complexes, SUMOylation prevents Rad52 from forming nucleolar foci (Torres-Rosell et al., 2007). Nucleolar Rad52 localization is further prevented by condensin-mediated rDNA condensation and RNA polymerase I transcription (Tsang and Zheng, 2009). Conversely, CLIP or cohibin disruption promotes rDNA release and aberrant recombination (Mekhail et al., 2008).

Keeping rDNA repeats within the nucleolus prevents aberrant recombination between different units; however, this spatial confinement presents a threat when cells encounter DNA damage. Damaged rDNA repeats need to be released from the nucleolus for repair by the HR machinery, and re-enter once fixed (Harding et al., 2015; van Sluis and McStay, 2015; Torres-Rosell et al., 2007). Similarly, repair of centromeric heterochromatin is blocked by Smc5/6-dependent exclusion of Rad51 in *Drosophila* and mouse cells, and its recovery involves the relocation of the damaged locus outside the heterochromatin domain (Chiolo et al., 2011; Tsouroula et al., 2016).

In addition to excluding Rad52 from the nucleolus, SUMO also mediates relocation of unresolved DSBs and damaged heterochromatin to the nuclear periphery for DNA repair (Horigome and Gasser, 2016). However, while the molecular events in the relocation of centromeric heterochromatin and other damaged sites have been intensely studied (Churikov et al., 2016; Horigome et al., 2016, 2019; Kalocsay et al., 2009; Ryu et al., 2015), the mechanisms controlling the relocation of broken rDNA repeats have remained mostly elusive. Here, we report the molecular events that regulate nucleolar rDNA release in *S. cerevisiae*. We find that a series of posttranslational modifications control association of key molecular players at the nuclear periphery. We show that blocking rDNA relocation using a constitutively bound CLIP-cohibin complex is detrimental for cell survival, even in the absence of external DNA damage. This result implies that CLIP-cohibin dissociation is dynamic. We find that Nur1 phosphorylation is a critical trigger for disruption of the CLIP-cohibin complex (hereafter referred to as rDNA tethering complex). In addition, our data reveal that SUMOylation of CLIP-cohibin mediates movement of individual rDNA repeats out of the nucleolus. SUMOylated CLIP-cohibin recruits the Cdc48/p97 segregase to assist in the disassembly of the rDNA tethering complex, a mechanism conserved in human cells. We propose that phosphorylation and SUMOylation of the CLIP-cohibin complex facilitate the transient release of damaged rDNA repeats from the nucleolus to the nucleoplasm to allow DNA repair.

## Results

### Permanent rDNA tethering is detrimental for cell survival

Deletion of CLIP or cohibin causes rDNA release from the nucleolus, which is reminiscent of rDNA relocation observed during DNA damage repair (Mekhail et al., 2008; Torres-Rosell et al., 2007). We thus postulated that rDNA release requires dynamic dissociation of CLIP and cohibin. To test this hypothesis, we generated an inducible artificial tethering system by fusing Heh1 (CLIP) to GFP-binding protein (GBP) under the control of the *GAL1* promoter and expressing it with ^GFP^Lrs4 (cohibin) (Figure 1a). Intriguingly, when induced by galactose, these fusion proteins caused a severe growth phenotype, implying that failure to dissociate the CLIP-cohibin complex is toxic for cells (Figure 1b, lane 4). A similar phenotype was observed when co-expressing GFP-fusions of the rDNA-bound proteins Net1 and Tof2 with Heh1-GBP (Fig 1a,b, lanes 6 and 8) (Huang et al., 2006). Expressing Heh1 directly fused to either Lrs4 or Tof2 also produced severe growth defects (Figure 1c, lanes 4 and 7). In contrast, perinuclear tethering of the unrelated genomic Gal4-DNA-binding domain did not affect growth (Figure 1c, lane 8). Notably, growth was fully restored by expression of linear Heh1-Lrs4 fusions with mutations preventing cohibin assembly (*lrs4 Δ1-35*) (Corbett et al., 2010) or impairing its interaction with downstream factors (*lrs4 Q325Stop*) (Johzuka and Horiuchi, 2009) (Figure 1c, lanes 5 and 6, respectively). We verified that all fusion variants displayed similar expression levels, excluding any lack of growth defect due to altered protein levels. The only exception was the expression of Heh1-Tof2, which was undetectable despite the substantial phenotype caused by its expression (Supplementary Figure 1b). Moreover, the observed phenotypes correlated with the expression levels, as driving Heh1-Lrs4 expression with the weaker *GALS* promoter resulted in less pronounce growth defects (Supplementary Figure 1c,d). Based on these observations, we conclude that cell viability critically depends on the dissociation of ribosomal repeats from the nuclear periphery and nucleolus, and dynamic CLIP-cohibin disassembly.

**Figure 1.**
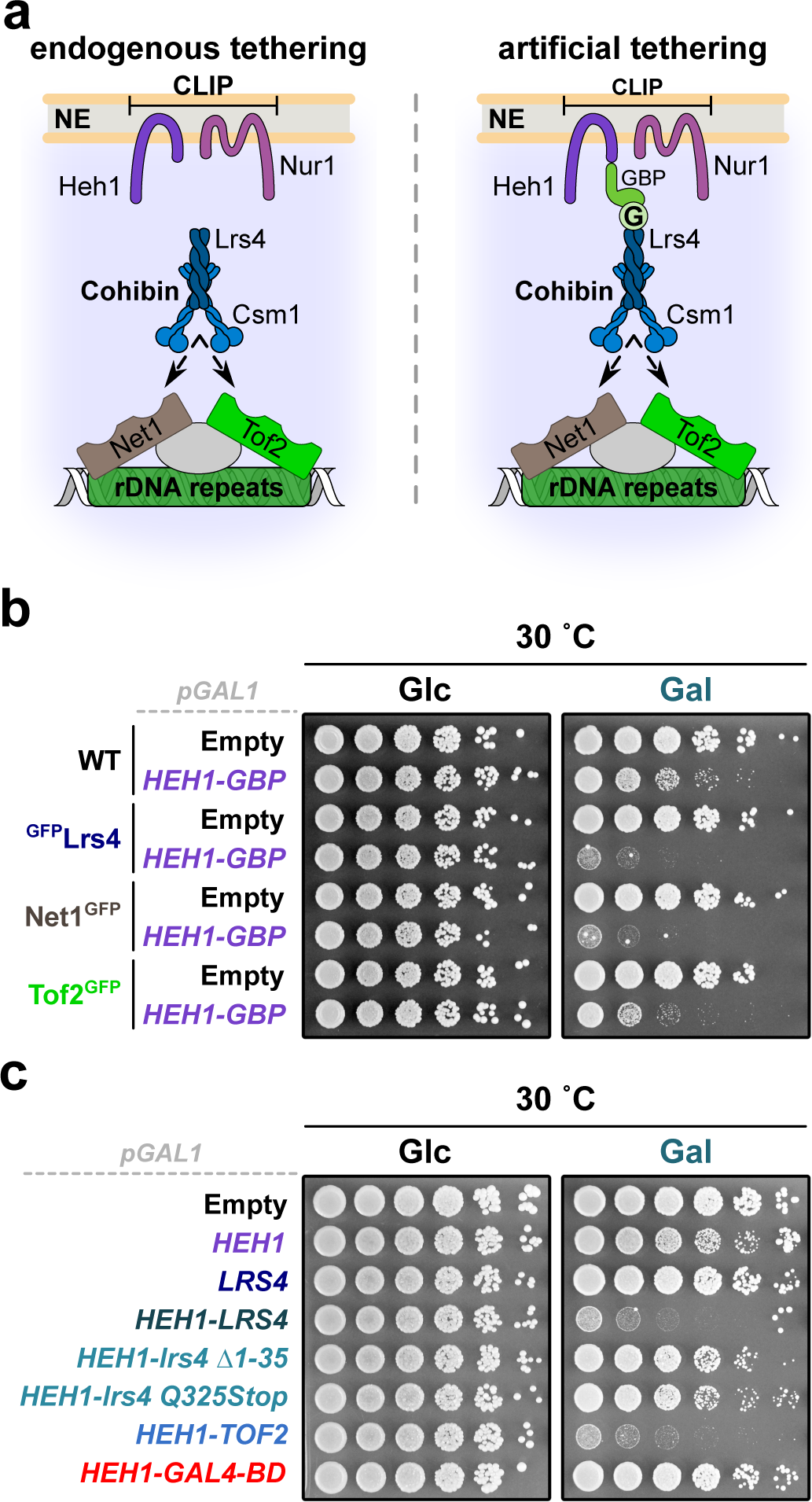
Constitutive perinuclear tethering of rDNA is lethal. **a**, Scheme of synthetic rDNA tethering through expression of GFP- and GBP-fusion proteins. Lrs4 binds ribosomal repeats through interaction with the rDNA-bound proteins Net1 or Tof2. G, GFP. **b**, Five-fold serial dilutions of WT, ^GFP^Lrs4, Net1^GFP^ and Tof2^GFP^ cells transformed with empty vector or a plasmid bearing *HEH1* fused to GFP-binding protein (GBP). **c**, Five-fold serial dilutions of WT cells transformed with empty vector or plasmids bearing the indicated GFP-tagged fusion proteins. For **b** and **c**, cells were spotted and grown on selective media with glucose (control, Glc) or galactose (induction, Gal) at 30°C for 3 days. All constructs contain the galactose-inducible promoter.

### The Nur1 C-terminal domain mediates rDNA tethering complex association

To unveil the molecular mechanism that controls CLIP-cohibin disassembly, we dissected interactions within the rDNA tethering complex. Using co-immunoprecipitation (coIP) assays, we found that deleting individual CLIP or cohibin components affects the binding of the two complexes. Specifically, Heh1 (CLIP) failed to interact with Csm1 (cohibin) in *lrs4Δ* or *nur1Δ* cells (Figure 2a,b). Moreover, Lrs4 (cohibin) interaction with Nur1 (CLIP) was abolished in cells lacking Csm1 (Supplementary Figure 2a). These results indicate that the integrity of both complexes is required for CLIP-cohibin association. In contrast, Heh1 interaction with Nur1 was largely unaffected in the absence of Lrs4 (Supplementary Figure 2b), revealing that CLIP formation is independent of cohibin integrity. In *S. cerevisiae*, *HEH1* is a rare example of alternative splicing, which generates a shorter version known as Heh1-S that does not bind Nur1 (Capella et al., 2020; Grund et al., 2008). We consistently detected Lrs4 binding to Heh1, but not Heh1-S, upon expression of an N-terminal GFP-Heh1 fusion that allows detection of both splice variants (Supplementary Figure 2c).

**Figure 2.**
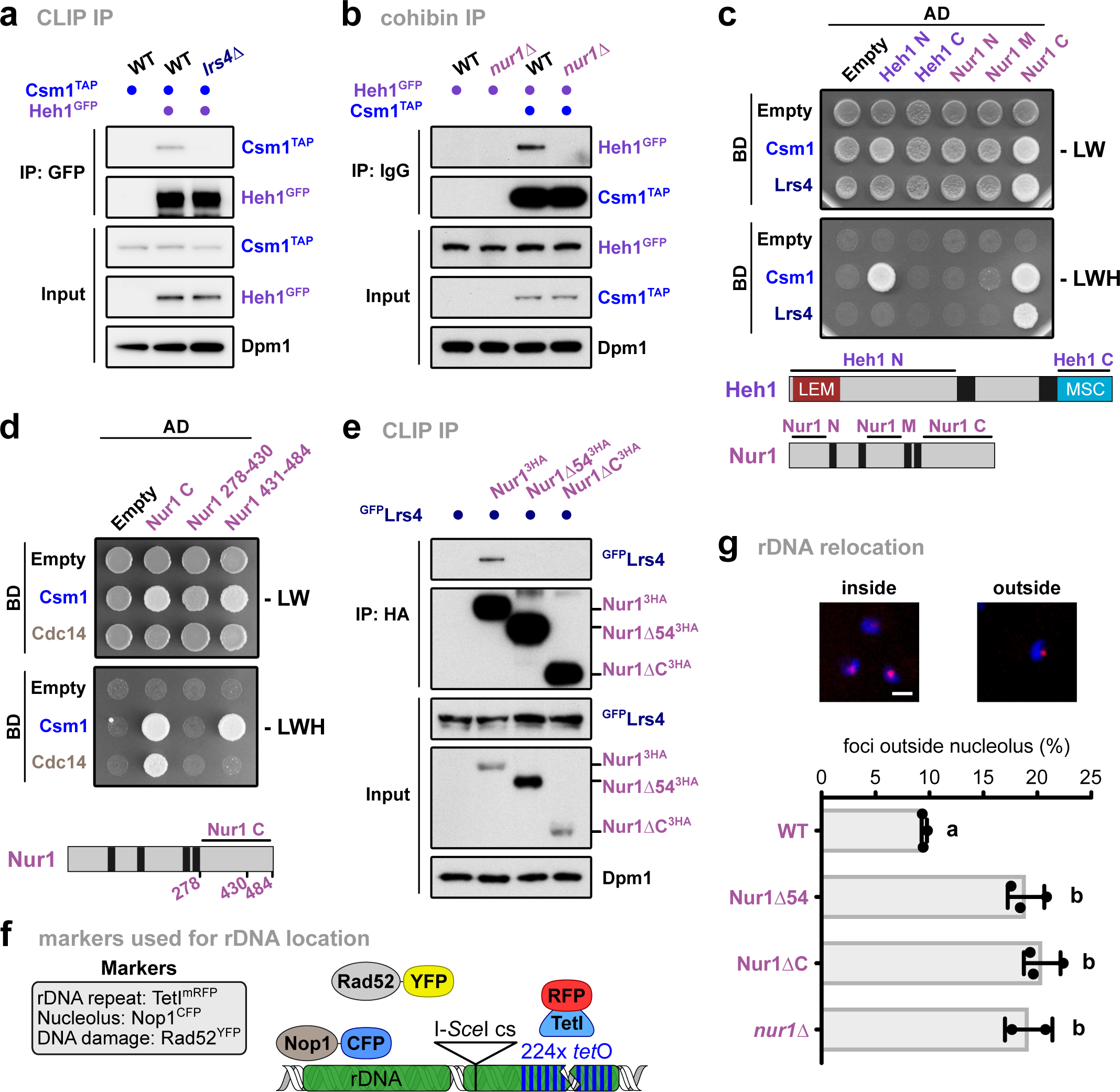
The Nur1 C-terminus is critical for CLIP-cohibin interaction. **a** and **b**, Co-immunoprecipitation of Csm1^TAP^ and Heh1^GFP^ in WT, *lrs4Δ* (**a**) and *nur1Δ* (**b**) cells. **c** and **d**, Y2H analysis of Csm1 and Lrs4 with Heh1 and Nur1 nucleoplasmic domains (**c**); or of Csm1 and Cdc14 with Nur1 C and truncations (**d**). Fusions with Gal4-activating domain (AD) or Gal4-DNA-binding domain (BD) are indicated. Cells were spotted on control media (-LW) or selective media (-LWH) and grown for 3 days. Scheme of the constructs used for CLIP complex components is shown. **e**, Co-immunoprecipitation of ^GFP^Lrs4 with either HA-tagged Nur1 full-length (Nur1^3HA^), lacking its last 54 residues (Nur1Δ54^3HA^) or the complete C-terminal domain (Nur1ΔC^3HA^). **f**, Scheme of rDNA locus and fluorescent markers used in **g**. Cells bear a tetO array adjacent to an I-SceI endonuclease cut site inserted into an rDNA unit on chromosome XII, which is revealed by TetImRFP foci. These cells also express Rad52^YFP^ as a marker of the HR machinery. The nucleolus is visualized by a plasmid expressing the nucleolar protein Nop1^ECFP^. **g**, Percentage of undamaged Nur1^6HA^, Nur1Δ54^6HA^, Nur1ΔC^6HA^ or *nur1Δ* cells with rDNA repeats localized outside the nucleolus. Repeat location was monitored by the position of TetImRFP focus relative to the nucleolar mark, and quantification is shown. Data are mean of n = 2-3 independent biological replicates. Representative images are shown. Scale bar, 2 µm. Analysis of variance (ANOVA) was performed, and different letters denote significant differences with a Tukey’s *post hoc* test at P < 0.05. For immunoblots, Dpm1 served as loading control.

Using yeast two-hybrid assays (Y2H), we mapped interactions between individual domains. We found that the N-terminal Heh1 domain is critical for Csm1 association, and the C-terminal Nur1 region interacts with both cohibin members (Figure 2c). Further analysis narrowed the Nur1 interaction interface with Csm1 down to the last 54 residues of the C-terminus (Figure 2d, Supplementary Figure 2d). Using coIP experiments, we confirmed the critical role of these residues for Nur1 association with Lrs4 *in vivo* (Figure 2e). Consistent with this, removal of the last residues also abolished Nur1 binding to Sir2 (Supplementary Figure 2e), which associates with Nur1 through cohibin (Huang et al., 2006). In contrast, deletion of these C-terminal Nur1 residues does not impact CLIP complex formation (Supplementary Figure 2e).

To monitor changes in the location of individual rDNA repeats, we used a yeast strain that harbors a tandem array of Tet-repressor-binding sites (224x*tetO*) inserted in the rDNA locus (Torres-Rosell et al., 2007). This strain also expresses TetI^mRFP^ to localize the modified rDNA unit as well as Rad52^YFP^ and Nop1^CFP^ to visualize Rad52 foci and the nucleolus, respectively (Figure 2f). Compared to WT cells, mutants lacking the Nur1 C-terminal domain or its last 54 residues displayed a marked increased in the number of cells with TetI^mRFP^ foci outside the nucleolus, reaching levels similar to *nur1Δ* cells (Figure 2g). From these data, we conclude that the most C-terminal region of Nur1 is required for CLIP-cohibin association.

### Phosphorylation of the CLIP complex disrupts rDNA tethering

Nur1 was shown to be phosphorylated (Godfrey et al., 2015), and this modification is removed by the nucleolar phosphatase Cdc14 recognizing the C-terminal domain (Figure 2d). As the C-terminal domain is also required for cohibin interaction, we tested whether phosphorylation triggers the disassembly of the rDNA tethering complex. Indeed, under conditions that favor the accumulation of phosphorylated Nur1, using either a thermosensitive mutant of Cdc14 (*cdc14-3*) or blocking globally dephosphorylation, we found, using coIP, that interaction of Nur1 with Lrs4 was reduced (Figure 3a, Supplementary Figure 3a). Conversely, depleting Net1, which inhibits Cdc14 phosphatase activity (Shou et al., 1999), increased Nur1 interaction with Lrs4 (Supplementary Figure 3b). To determine target sites in Nur1, we mutated several residues known to be phosphorylated (Holt et al., 2009) to aspartic acid (D) mimicking the modified state (Supplementary Figure 3c). Notably, using Y2H assays, we found that the Nur1 interaction with Csm1 was abolished when residues S441, T446 and S449 were mutated to D, but unaffected when replaced by alanine (Supplementary Figure 3d). We confirmed the critical role of these residues for the CLIP-cohibin interaction *in vivo* using coIP (Figure 3b).

**Figure 3.**
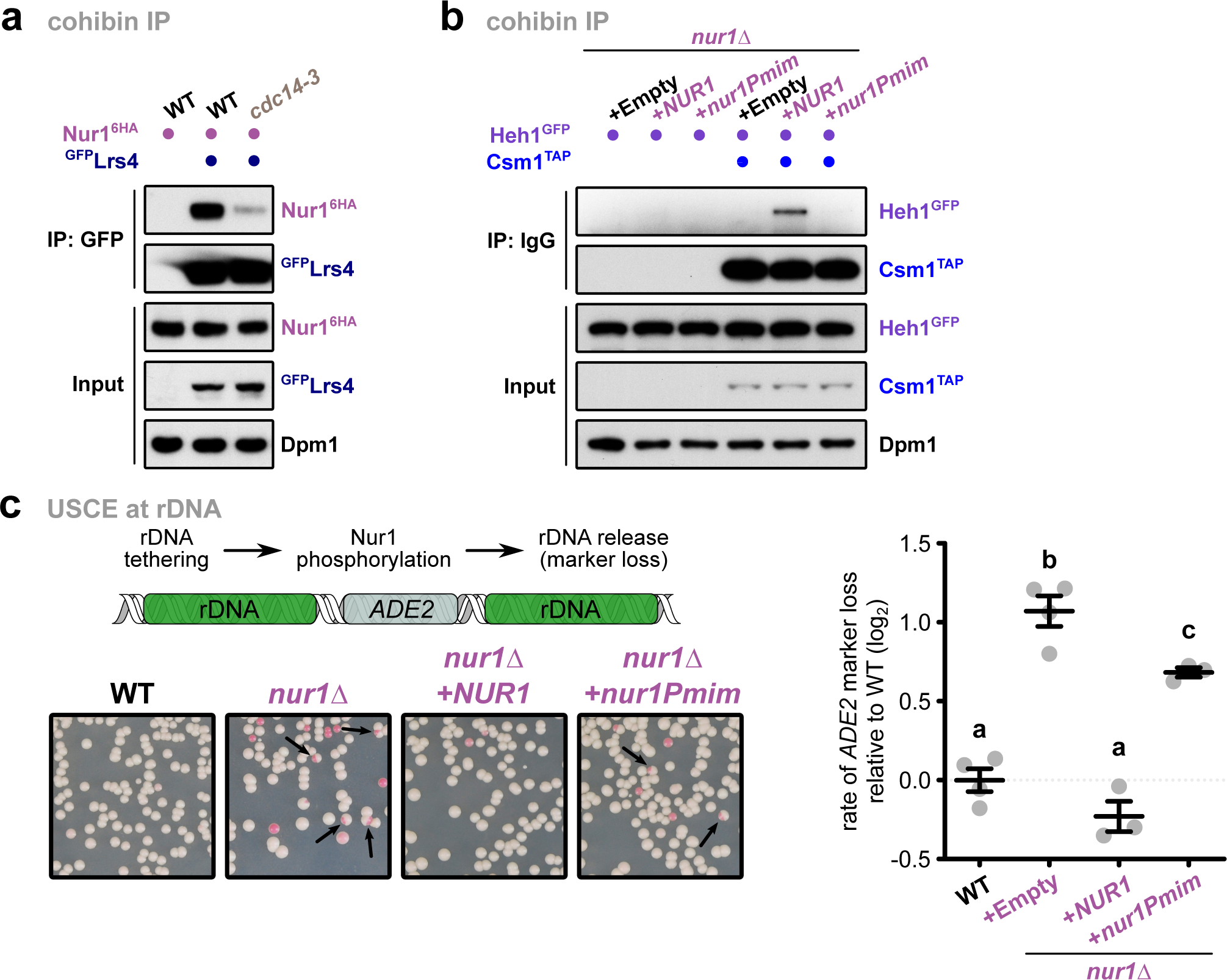
C-terminal Nur1 phosphorylation disrupts the rDNA tethering complex. **a**, Co-immunoprecipitation of Nur1^6HA^ with ^GFP^Lrs4 in WT or *cdc14-3* mutant cells. Cells were grown at the permissive temperature and shifted to 37°C for 1h. **b**,Co-immunoprecipitation of Heh1^GFP^ with Csm1^TAP^ in *nur1Δ* cells. The strains were transformed with empty vector or plasmids bearing *NUR1* or its phosphomimetic mutant (*nur1Pmim*) expressed from the endogenous promoter. **c**, Quantification of rDNA recombination rates in WT or *nur1Δ* cells as measured by unequal sister chromatid exchange (USCE) using the *ADE2* marker inserted into rDNA. Cells have been transformed with empty vector or plasmids bearing *NUR1* or *nur1Pmim* expressed from the endogenous promoter. The rate of marker loss is calculated as the ratio of half-sectored colonies (as indicated by the arrows) to the total number of colonies, excluding completely red colonies, shown in log_2_ scale relative to WT. Data are mean of n = 3-4 independent biological replicates. ANOVA was performed, and different letters denote significant differences with a Tukey’s *post hoc* test at P < 0.05. For immunoblots, Dpm1 served as loading control.

Loss of perinuclear tethering causes destabilization of rDNA repeats, which can be quantitated by unequal sister-chromatid exchange (USCE) using the *ADE2* reporter gene inserted into the rDNA locus. When assessing the rate of *ADE2* gene loss in the *nur1* phosphomimic mutant, we found increased USCE similar to the *nur1Δ* mutant (Mekhail et al., 2008). This indicates that Nur1 phosphorylation decreases rDNA stability (Figure 3c). These results imply that the CLIP-cohibin association is dynamically controlled by Nur1 phosphorylation and counterbalanced by the phosphatase Cdc14.

### CLIP-cohibin dissociation depends on SUMOylation

SUMOylation plays a prominent role in the DNA damage response and rDNA stability (Jalal et al., 2017). As the relocation of other genomic regions after damage requires SUMO (Horigome and Gasser, 2016), we examined whether this modification also facilitates rDNA release from the nucleolus. Testing the CLIP-cohibin interaction, we found that SUMO overexpression decreased Nur1 association with Lrs4 but not with its partner Heh1 (Figure 4a, Supplementary Figure S4a). We further found that SUMO overexpression results in high *ADE2* marker loss, indicating decreased rDNA stability using USCE (Figure 4b). Notably, additional deletion of *LRS4* resulted in a non-additive phenotype, suggesting that SUMOylation and CLIP-cohibin act within the same pathway (Figure 4b). Consistently, cells expressing the SUMO isopeptidase mutant *ulp2ΔC,* which lacks the region required for Csm1 interaction (Liang et al., 2017), also showed an increased rate of marker loss (Supplementary Figure 4b). When studying the location of rDNA using an mRFP-marked repeat insertion, SUMO overexpression increased the number of cells with delocalized rDNA repeats (Figure 4c), further supporting the role of SUMO in promoting rDNA release.

**Figure 4.**
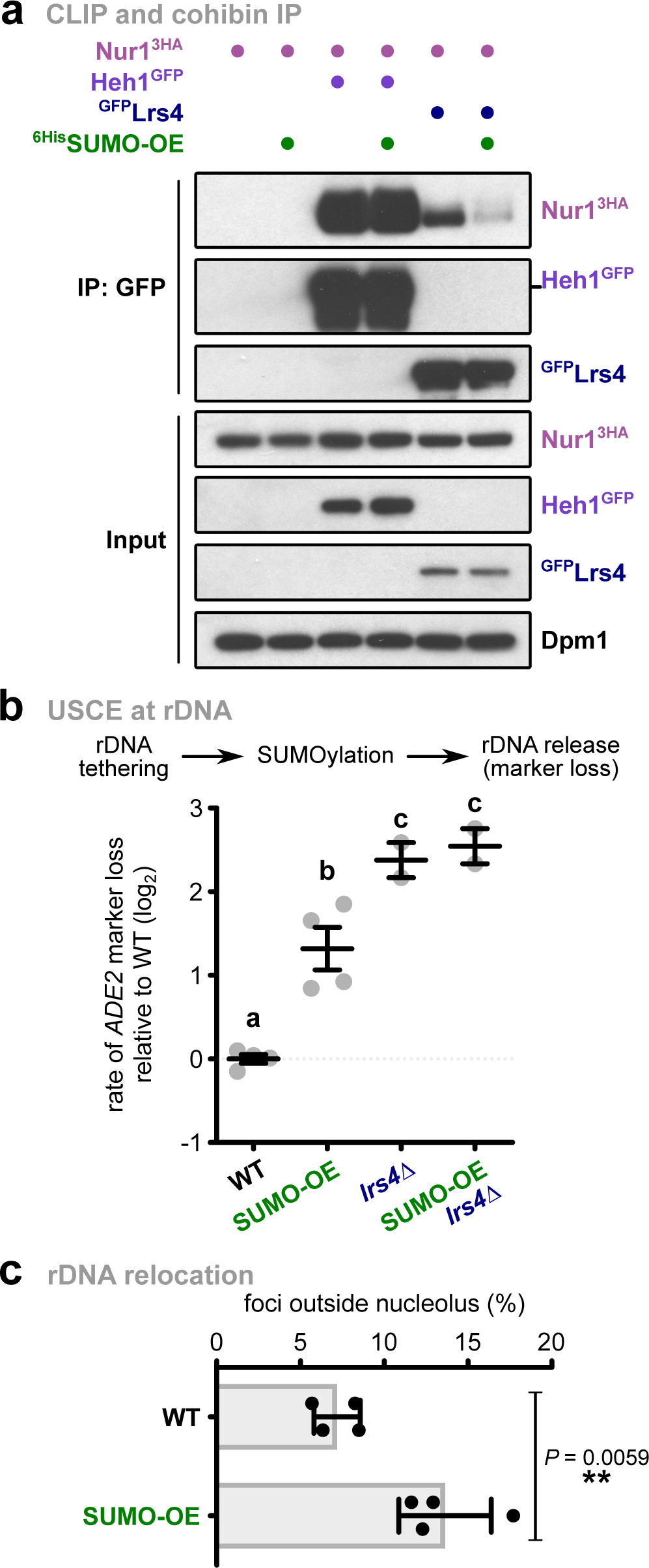
SUMO overexpression promotes CLIP-cohibin disassembly and rDNA relocation and recombination. **a**, Co-immunoprecipitation of Nur1^3HA^ with Heh1^GFP^ or ^GFP^Lrs4 in cells expressing SUMO at endogenous levels or overexpressed from the *ADH1* promoter (^6His^SUMO-OE). Dpm1 served as loading control. **b**, Rates of unequal rDNA recombination in WT or *lrs4Δ* cells expressing SUMO at endogenous levels or overexpressed from vectors with the *GAL1* promoter (SUMO-OE), grown in galactose-containing media for 4 days. Data are mean of n = 4 independent biological replicates for WT cells, and n = 2 for *lrs4Δ* strains, shown in log_2_ scale relative to WT. Marker loss is calculated as in Figure 3. **c**, Percentage of undamaged cells with endogenous (WT) or overexpressed (SUMO-OE) SUMO levels with rDNA repeats localized outside the nucleolus, monitored as described for Figure 2. Quantification of the marked rDNA unit was scored from 4 independent biological replicates, and the mean is shown. Analysis of variance was performed, and different letters denote significant differences with a Tukey’s *post hoc* test at P < 0.05.

In contrast, transcript levels of the non-transcribed spacers NTS1 and NTS2 were not altered, indicating that the SUMO overexpression-mediated release of ribosomal repeats does not decrease rDNA silencing (Supplementary Figure 4c), which is also consistent with previous reports in CLIP deficient cells (Mekhail et al., 2008). Furthermore, Csm1 association with NTS1 was not reduced, but instead enhanced under these conditions (Supplementary Figure 4d). These data suggest that cohibin remains bound to rDNA after CLIP dissociation.

Since both SUMOylation and Nur1 phosphorylation stimulate CLIP-cohibin dissociation, we tested whether SUMO overexpression induces Nur1 phosphorylation. To this end, we monitored the electrophoretic mobility of Nur1 using Phos-tag gels in cells with endogenous or elevated levels of SUMO. Nur1 phosphorylation levels change in a cell-cycle-dependent manner, in agreement with previous reports (Godfrey et al., 2015), and SUMO overexpression did not affect these changes (Supplementary Figure 4e). This result indicates that Nur1 phosphorylation occurs independently of SUMOylation, either upstream or in parallel. Overall, these findings argue that SUMOylation disrupts the CLIP-cohibin complex, promoting the release of ribosomal repeats.

### Multiple members of the rDNA tethering complex are SUMOylated

To obtain further insights into the mechanism triggering SUMO-dependent disruption of the CLIP-cohibin complex, we used a candidate Y2H approach to identify potential SUMO substrates. Both covalent and/or non-covalent SUMO association with its targets can result in the reconstitution of the Gal4 transcription factor in Y2H assays. To distinguish between these interaction modes, we used a SUMO conjugation-deficient mutant (SUMO^AA^) (Höpfler et al., 2019; Pfander et al., 2005) to test binding of partners. Using this approach, we found that all CLIP-cohibin members interact with SUMO but not SUMO^AA^ (Supplementary Figure 5a), suggesting that the rDNA tethering complex is subject to SUMOylation.

To validate these results, we examined Nur1 SUMOylation *in vivo*. Upon SUMO overexpression, Nur1 immunoprecipitation revealed an additional, slower migrating form that was further shifted when SUMO was fused to GFP (Figure 5a). These modifications were absent in cells lacking the SUMO E3 ligase Siz2 (Figure 5a). We further confirmed Nur1 SUMOylation by enriching for ^6His^SUMO conjugates using Ni-NTA pulldowns under denaturing conditions (Figure 5b). Interestingly, SUMOylated Nur1 was also reduced in *lrs4Δ* cells and *sir2Δ* cells, or when cells were treated with nicotinamide (Figure 5b, Supplementary Figure 5b,c), a strong inhibitor of Sir2 activity (Bitterman et al., 2002). Together, these results imply that Nur1 is SUMOylated in a Siz2-dependent manner, and this requires CLIP association with the rDNA locus. When testing cohibin members, we failed to detect SUMOylated Lrs4 among ^6His^SUMO conjugates using Ni-NTA pulldowns (data not shown). Therefore, we overexpressed an isopeptidase-resistant mutant of SUMO (SUMO^Q95P^) to increase the sensitivity of SUMO conjugate detection(Pabst et al., 2019). Immunoprecipitation of Lrs4 under semi-denaturing conditions, which remove non-covalent interactions (see methods), revealed an additional slower-migrating band recognized by an anti-SUMO antibody (Figure 5c). We concluded that multiple members of the CLIP-cohibin complex are SUMOylated *in vivo*.

**Figure 5.**
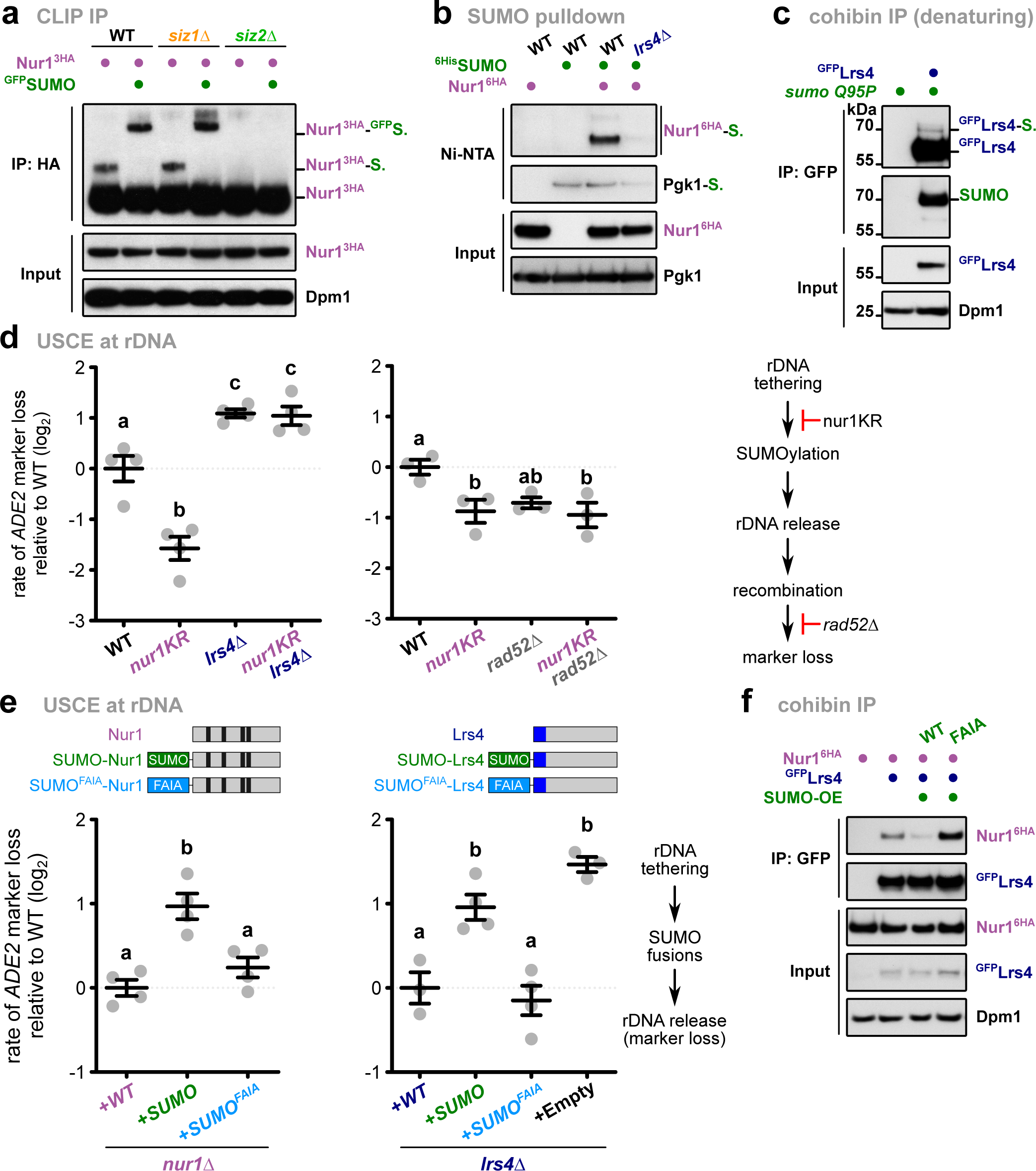
SUMOylation of Nur1 and Lrs4 triggers CLIP-cohibin dissociation. **a**, Immunoprecipitation of Nur1^3HA^ in WT, *siz1Δ* and *siz2Δ* expressing SUMO (endogenous promoter) or ^GFP^SUMO (*ADH1* promoter). Bands corresponding to Nur1 unmodified or monoSUMOylated are labeled. **b**, Denaturing Ni-NTA pulldowns of ^6His^SUMO conjugates from WT or *lrs4Δ* cells expressing and Nur1^6HA^, as indicated. Pgk1 SUMOylation was analyzed to control for pulldown efficiency, while unmodified Pgk1 served as loading control. **c**, Immunoprecipitation under semi-denaturing conditions of ^GFP^Lrs4 in cells overexpressing an isopeptidase-resistant SUMO mutant (SUMO^Q95P^). Detection of SUMO from immunoprecipitated samples is shown. **d**, Rates of unequal rDNA marker loss in WT, *lrs4Δ* (left) or *rad52Δ* (right) cells expressing endogenously 6HA-tagged Nur1 or the SUMOylation deficient Nur1 K175-176R mutant (*nur1KR*). **e**, Rates of unequal rDNA marker loss in *nur1Δ* or *lrs4Δ* cells transformed with empty vector or plasmids bearing 3HA-tagged *NUR1* (left), TAP-tagged *LRS4* (right) or the indicated linear fusions with the endogenous promoter, as indicated. Used constructs for Nur1 and Lrs4 are shown. Black, predicted transmembrane domains; blue: Csm1-interacting region. **f**, Co-immunoprecipitation of Nur1^6HA^ with ^GFP^Lrs4 in cells with endogenous SUMO levels, or strains overexpressing either SUMO WT or a mutant unable to recognize SIMs (FAIA, which harbors the mutations F37A I39A). The different strains were grown to mid-log phase, and SUMO variants were induced by adding 2% galactose for 2 h 30’. For **a**, **c** and **f**, Dpm1 served as loading control. For **d** and **e**, the rate of marker loss is calculated as in Figure 3; data are the mean of n = 3-4 independent biological replicates, shown in log2 scale relative to WT. ANOVA was performed, and different letters denote significant differences with a Tukey’s post hoc test at P < 0.05.

To examine whether SUMOylation of the CLIP-cohibin complex affects rDNA stability, we determined the SUMO target sites on Nur1 by mutating putative acceptor lysine residues (K) to arginine (R). Using Ni-NTA pulldowns with ^6His^SUMO and IP assays, we found that combined mutation of the acceptor lysine residues K175 and K176 abolished SUMOylation of Nur1 (Supplementary Figure 5d,e). Notably, replacing Nur1 with the SUMO-deficient mutant (*nur1KR*) resulted in decreased rDNA marker loss, which was entirely dependent on the presence of Lrs4 (Figure 5d, left). Moreover, while *RAD52* deletion decreases USCE as previously reported (Torres-Rosell et al., 2007), additional mutation of SUMOylation acceptor sites in Nur1 did not further reduce USCE in *rad52Δ* (Fig 5d, right). Overexpression of SUMO partially alleviates the USCE phenotype of *nur1KR* (Supplementary Figure 5f, left). To further validate the effect of SUMO on rDNA stability, we generated strains expressing Nur1 fused to the SUMO protease domain of Ulp1 (Nur1-UD), which is expected to reduce SUMOylation of the rDNA tethering complex (Colomina et al., 2017). Notably, the presence of Nur1-UD decreased USCE levels compared to the fusion with the catalytically dead domain of Ulp1 (Nur1-uD^i^; Supplementary Figure 5f, right). Together, this argues that Nur1 SUMOylation acts upstream of Rad52-mediated recombination, and that increased SUMOylation of other CLIP-cohibin members can compensate for the loss of Nur1 SUMOylation.

To validate the role of SUMOylation in the rDNA tethering complex through a gain-of-function approach, we expressed Nur1 and Lrs4 as linear fusions with SUMO in *nur1Δ* and *lrs4Δ* cells, respectively, and examined rDNA stability by USCE. Notably, while expressing non-fused Nur1 and Lrs4 rescued the deletion mutants, the SUMO fusion proteins failed to complement the USCE phenotype (Figure 5e). Several proteins have been shown to interact with SUMO with moderate affinity through their SIMs (Geiss-Friedlander and Melchior, 2007). We therefore introduced mutations into the SUMO moiety preventing its recognition by SIMs (SUMO^FAIA^; Supplementary Figure 5g,h) in the linear Nur1 and Lsr4 SUMO fusion proteins. When assessing rDNA recombination, these SUMO^FAIA^ fusions displayed USCE rates that are significantly lower than the corresponding WT-SUMO fusions, and comparable to the non-fusion proteins (Figure 5e). This implies that the specific recognition of SUMO in the rDNA tethering complex triggers rDNA recombination. In agreement with this observation, we found that the Lrs4 interaction with CLIP is strongly reduced when WT SUMO is overexpressed, but not the SIM mutant (Figure 5f). These data argue that SUMOylation on CLIP-cohibin, likely through SUMO-dependent recruitment of other protein(s), promotes rDNA relocation.

### Cdc48 recognizes the CLIP-cohibin complex and supports rDNA release

The AAA-ATPase Cdc48 (also known as VCP/p97) acts as a segregase to disassemble protein complexes during proteasomal degradation, but also has functions in non-proteolytic pathways, such as DNA damage repair (Franz et al., 2016). Besides recognizing ubiquitylated proteins, Cdc48 has also been reported to bind to SUMOylated proteins (Bergink et al., 2013; Nie et al., 2012). Therefore, we tested whether the CLIP-cohibin complex is targeted by Cdc48 in a SUMO-dependent manner. We found that Cdc48 interacts weakly with Lrs4 in WT cells using coIP. This interaction was increased in cells overexpressing either a thermosensitive allele (*cdc48-6*) or an ATPase deficient mutant (*cdc48 E588Q*; Figure 6a), known to block or delay the release of substrates, respectively (Bergink et al., 2013; Bodnar and Rapoport, 2017). We also observed increased Cdc48 interaction with Nur1 in *cdc48-6* cells (Figure 6b). When examining the CLIP-cohibin association, Lrs4 binding to Nur1 was modestly increased in *cdc48-6* or *cdc48 E588Q* cells (Figure 6c). Together, these results suggest that Cdc48 assists in the disruption of the rDNA tethering complex.

**Figure 6.**
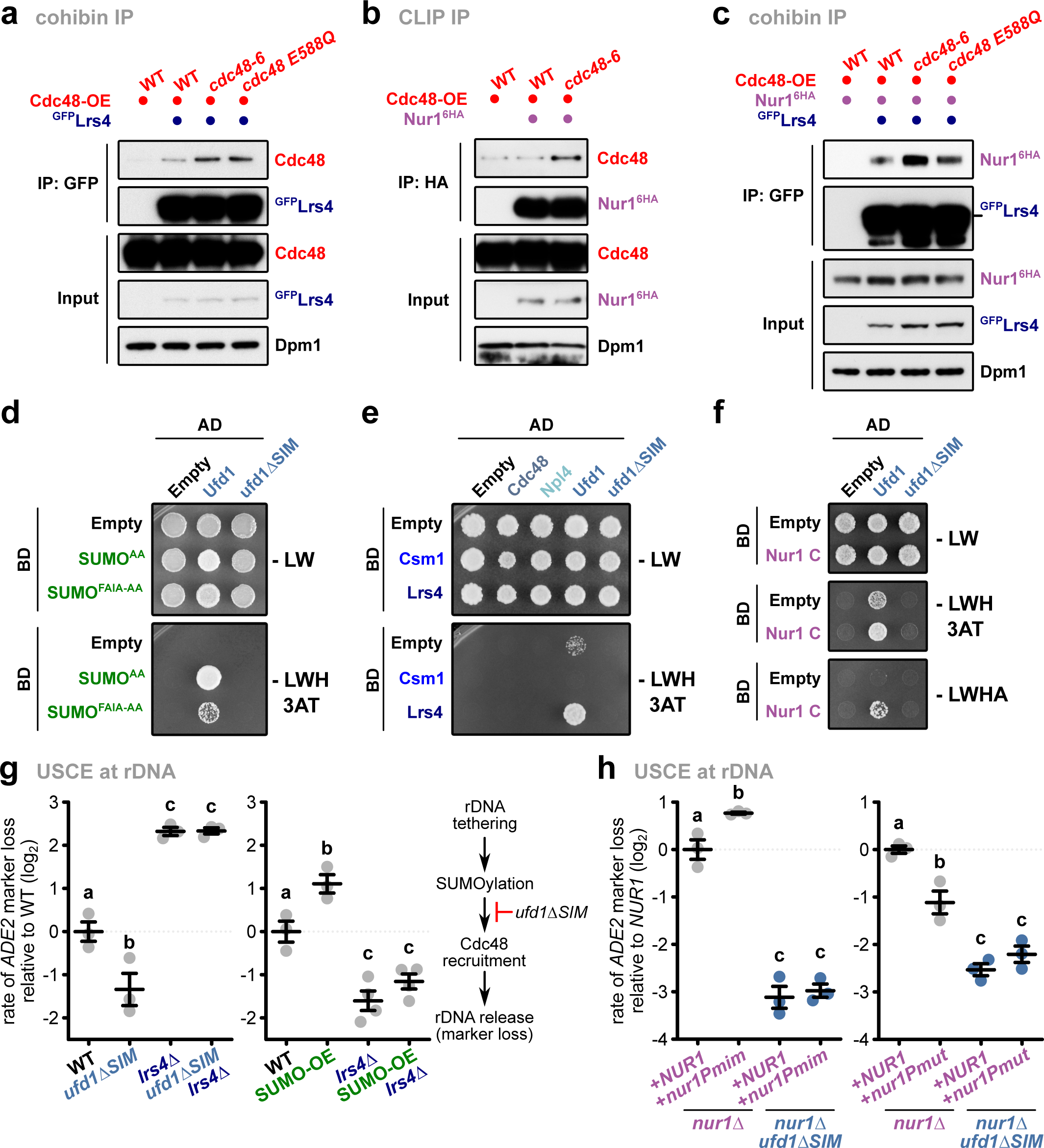
SUMOylation recruits Cdc48 via its co-factor Ufd1 to assist in rDNA release. **a** and **b**, Co-immunoprecipitation of Cdc48 with ^GFP^Lrs4 (**a**) or Nur1^6HA^ (**b**) in cells overexpressing WT or mutant Cdc48. **c**, Co-immunoprecipitation of Nur1^6HA^ with ^GFP^Lrs4 in cells overexpressing WT or mutant Cdc48. **d**, Y2H analysis of conjugation-deficient SUMO (SUMO^AA^) or the F37A I39A mutant unable to recognize SIMs (SUMO^FAIA-AA^) (as a Gal4-DNA-binding domain fusion, BD) with Ufd1 and ufd1ΔSIM (as a Gal4-activating domain fusion, AD). **e**, Y2H analysis of Csm1 and Lrs4 with Cdc48, Npl4, Ufd1 and ufd1ΔSIM. Fusions with Gal4-activating domain (AD) or Gal4-DNA-binding domain (BD) are indicated. **f**, Y2H analysis of Nur1 C (BD) with Ufd1 and ufd1ΔSIM (AD), as indicated. **g**, Rates of unequal rDNA recombination in WT, *ufd1ΔSIM*, *lrs4Δ* or *ufd1ΔSIM lrs4Δ* cells (left), in WT and *ufd1ΔSIM* strains expressing SUMO at endogenous levels or overexpressed from vectors with the *TEF1* promoter (SUMO-OE) (right). **h**, Rates of unequal rDNA recombination in nur1Δ and nur1Δ ufd1ΔSIM cells transformed with plasmids bearing NUR1 or its phosphomutant version (nur1Pmut) with the GAL1 promoter. Cells were grown in galactose-containing media for 4 days. For **a** and **c**, the different strains were grown to mid-log phase, and Cdc48 variants were induced by adding 2% galactose for 2 h 30’; Dpm1 served as loading control. For **d**, **e** and **f**, cells were spotted on control media (-LW) or selective media (-LWH + 3AT 1 mM) and grown for 3 days, or spotted on - LWHA selective media and grown for 4 days (**f**). For **g** and **h**, the rate of marker loss is calculated as in Figure 3; data are mean of n = 3-4 independent biological replicates, shown in log_2_ scale relative to WT (**g**) or *nur1Δ+NUR1* (**h**). ANOVA test was used, and different letters denote significant differences with a Tukey’s post hoc test at P < 0.05.

Cdc48 recognize SUMOylated proteins through its substrate-recruiting cofactor Ufd1 (Bergink et al., 2013; Nie et al., 2012) that contains a SIM motif (Figure 6d). Consistent with this, we found that both Lrs4 and Nur1 interact with Ufd1 in Y2H assays, and this interaction is SIM-dependent (Figure 6e,f). Notably, *ufd1ΔSIM* cells displayed a significant reduction of USCE, suggesting that recognition of SUMOylated Nur1 or Lrs4 is critical for rDNA release and recombination. In agreement with this notion, the decrease in USCE in *ufd1ΔSIM* mutant cells was completely relieved when Lrs4 was absent (Figure 6g, left). Furthermore, SUMO overexpression did not significantly alter the USCE phenotype of *ufd1ΔSIM* cells (Figure 6g, right), implying that Ufd1 acts downstream of SUMOylation in the maintenance of rDNA stability. We further tested the role of STUbLs, which have been reported to contribute to rDNA stability and recruit Cdc48 in a SUMO-dependent manner (Cal-Bkowska et al., 2011; Gillies et al., 2016; Liang et al., 2017). However, deletion of the STUbLs *SLX5*, *SLX8* or *ULS1* did not affect CLIP-cohibin association (Supplementary Figure 6). Together, these results suggest that Cdc48 recognizes the CLIP-cohibin complex, and promotes rDNA relocation in a SUMO- and Ufd1-dependent manner.

While our findings suggested that Nur1 phosphorylation does not depend on SUMOylation, both modifications may contribute to rDNA release through the same pathway, with SUMOylation/Cdc48 acting downstream of Nur1 phosphorylation. We therefore assessed how the phosphorylated states of Nur1 affects USCE in cells deficient in SUMO recognition. Notably, while expressing the phosphomimetic *nur1Pmim* mutant promoted rDNA recombination, this increase in USCE was blocked when *nur1Pmim* was combined with *ufd1ΔSIM* (Figure 6h, left). Conversely, cells deficient in Nur1 phosphorylation (*nur1Pmut*) that show reduced USCE rates displayed a non-additive phenotype in combination with the *ufd1ΔSIM* mutation (Figure 6h, right). Together, these results suggest that SUMOylation acts downstream of Nur1 phosphorylation.

### DNA damage triggers the dissociation of the rDNA tethering complex

DNA damage repair requires nuclear membrane release prior to the relocation of the rDNA repeats. To examine whether CLIP-cohibin disassembly plays a role in this physiological context, we studied its association and modification in response to double-strand breaks (DSBs). To this end, we transiently expressed the *Physarum polycephalum* homing endonuclease I-*Ppo*I, which generates DSBs at every rDNA repeat (Monnat et al., 1999; Muscarella and Vogt, 1993), from the galactose-inducible strong *GAL1* and moderate *GALL* promoters. Quantitative PCR analysis of genomic DNA confirmed that I-*Ppo*I expression induces DSBs within rDNA repeats in a dose-dependent manner without affecting the rDNA copy number (Figure 7a,b). When studying CLIP association with cohibin by coIP, we found that Heh1 binding to Lrs4 was reduced upon I-*Ppo*I induction (Figure 7a), whereas Lrs4 association with rDNA was not altered (Figure 7b), suggesting that rDNA breaks trigger CLIP-cohibin disassembly. In agreement with this finding, we observed that I-*Ppo*I induction increases Nur1 phosphorylation (Supplementary Figure 7a) and the appearance of global SUMO conjugates (Supplementary Figure 7b). To investigate whether preventing CLIP-cohibin disassembly affects rDNA release after damage, we performed live-cell imaging to monitor the nucleolar location of a *tetO*/TetI^mRFP^-marked repeat flanked by an I-*Sce*I endonuclease cut site. After I-*Sce*I induction, generation of a single DSB caused rDNA relocation to an extranucleolar site (Figure 7c), as previously reported (Torres-Rosell et al., 2007). Notably, rDNA relocation was substantially reduced in a *ufd1ΔSIM* mutant, which impairs SUMO recognition by Cdc48 and likely acts downstream of CLIP and cohibin SUMOylation (Figure 7c). Based on these results, we conclude that DSBs at the rDNA repeats promote nucleolar release through controlled disassembly of the tethering complex through Nur1 phosphorylation and SUMO-dependent Cdc48 activity.

**Figure 7.**
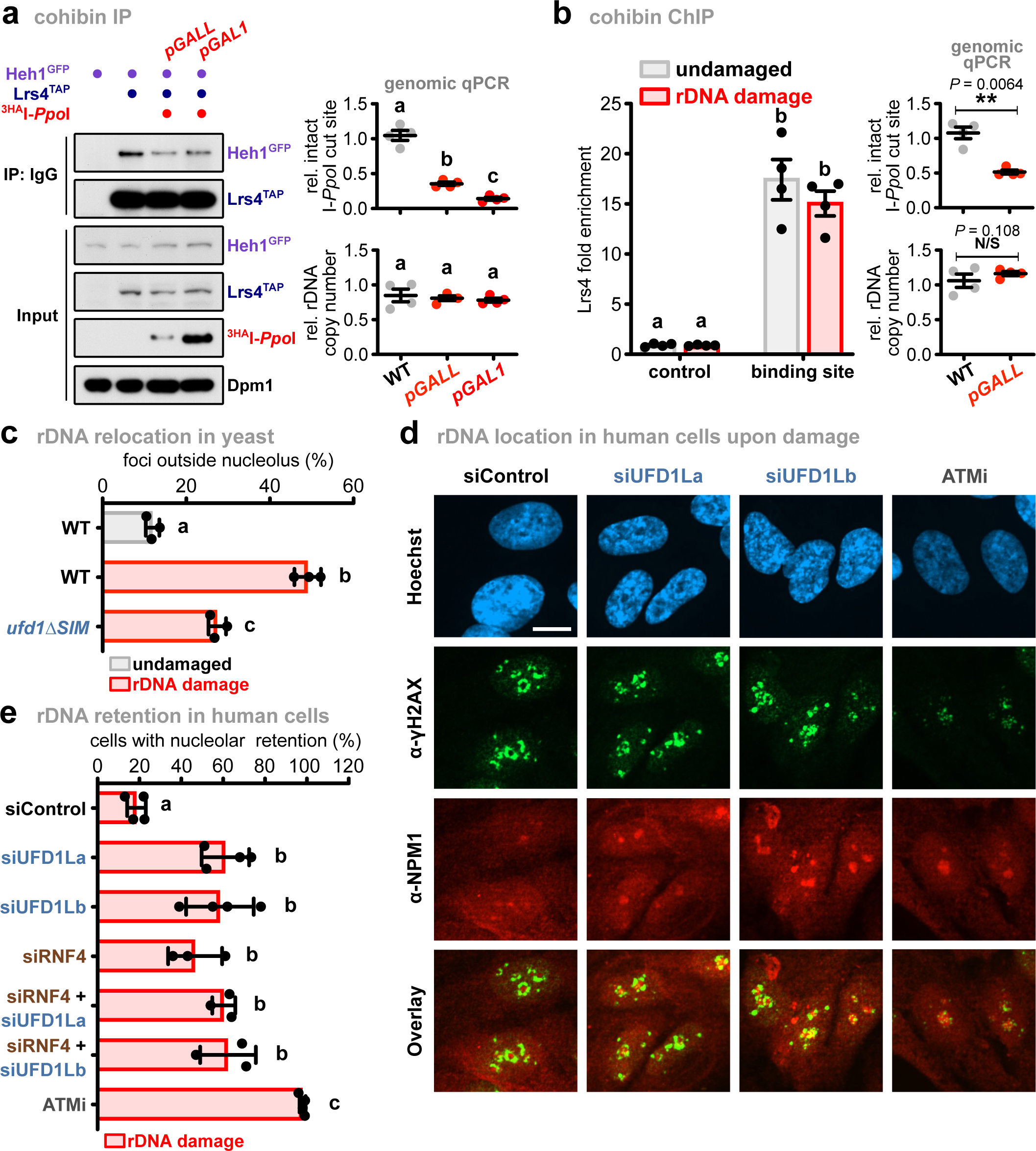
DNA damage contributes to CLIP-cohibin disassembly and Cdc48/p97-dependent rDNA relocation. **a**, Co-immunoprecipitation of Heh1^GFP^ with Lrs4^TAP^ in cells with or without rDNA damage. Dpm1 served as loading control. ^3HA^I-*Ppo*I with the galactose-inducible promoters *GALL* and *GAL1* were integrated at the *LEU2* locus. **b**, Lrs4^TAP^ binding to rDNA repeats in cells with or without rDNA damage, quantified by ChIP-qPCR. The ChIP values are shown as Lrs4 fold enrichment over the average of three rDNA positions, after normalization to input. Illustration shows a schematic representation of an rDNA unit. Amplified regions are highlighted with red arrows. **c**, Percentage of WT and *ufd1ΔSIM* cells with rDNA repeats localized outside the nucleolus before and after DSB induction. Relative nucleolar location of rDNA repeats was monitored and quantified as described for Figure 2. **d**, Retention of rDNA locus (NPM1) in human cells upon DNA damage treated with siRNA against UFD1L. Human RPE cells stably expressing FKBP12-^HA^I-*Ppo*I were transfected with siRNAs against UFD1L (siUFD1La and siUFD1Lb), or treated with the ATM kinase inhibitor KU55933 (ATMi). Representative images of immunofluorescence staining after rDNA damage, using the indicated antibodies, are shown. Scale bar, 10 µm. **e**, Quantification of human RPE cells transfected with siRNA against RNF4 (siRNF4), siUFD1L (siUFD1La and siUFD1Lb), or combination thereof, showing the percentage of cells with nucleolar retention of γH2AX foci, defined as cells that show an overlap (yellow) of the γH2AX (green) with the NPM (red) signal. For **a** and **b**, quantification of double-strand break (DSB) induction (top right) and rDNA copy number (bottom right) relative to undamaged cells is shown. For **b**, **c** and **e**, data are mean of n = 3-4 biologically independent experiments. ANOVA was performed, and different letters denote significant differences with a Tukey’s *post hoc* test at P < 0.05. N/S, not significant.

### UFD1 promotes rDNA release in human cells upon DNA damage

Since relocation of ribosomal repeats during DNA damage repair is a conserved mechanism (Harding et al., 2015; van Sluis and McStay, 2015), we speculated that key processes regulating rDNA dynamics might also be present in humans. To test this, we examined rDNA release upon damage in RPE (<u>r</u>etinal <u>p</u>igment <u>e</u>pithelium) cells depleted of the Ufd1 homolog, UFD1L. To generate DSBs in rDNA repeats, we used a tetracyclin-inducible expression system consisting of I-*Ppo*I fused to the FKBP12 destabilization domain, which can be stabilized by the addition of the compound shield-1 (Warmerdam et al., 2016). The movement of damaged repeats outside the nucleolus was assessed by quantifying the location of γH2AX foci, a marker for damaged DNA sites, relative to the nucleolar marker nucleophosmin (NPM). As previously reported (Harding et al., 2015; van Sluis and McStay, 2015; Warmerdam et al., 2016), broken repeats were released to a large extent from the nucleolus of control cells upon expression of I-*Ppo*I, as calculated by the number of cells with γH2AX foci within the nucleolus (Figure 7d,e). Upon depletion of UFD1L, we found that nucleolar segregation was reduced by more than 50% after damage relative to inhibition of the DNA damage checkpoint kinase ATM (<u>a</u>taxia telangiectasia-<u>m</u>utated), which resulted in a complete block of nucleolar release (Figure 7d,e, Supplementary Figure 7c). In contrast to yeast, the interaction between p97 and SUMO has not been described in humans. However, p97 is targeted to SUMOylated substrates with the aid of the STUbL RNF4 (Franz et al., 2016; Gibbs-Seymour et al., 2015; Sun et al., 2007). Notably, depletion of RNF4 blocks rDNA relocation upon damage, and additional UFD1L knock-down resulted in a non-additive phenotype, suggesting that RNF4 and p97 together assist relocation away from the nucleolus (Figure 7e). These results suggest that the role of Cdc48/p97 in controlling rDNA release during rDNA damage repair is conserved from yeast to humans.

## Discussion

The nucleolus shields the rDNA locus from uncontrolled recombination in eukaryotes, but this requires relocation to the nucleoplasm should DNA damage repair be needed (van Sluis and McStay, 2015; Torres-Rosell et al., 2007). However, how individual rDNA repeats exit the nucleolus was previously poorly understood. Here, we have identified the critical steps triggering the nucleolar release of rDNA repeats in yeast both under normal growth conditions and after exogenous DNA damage. These molecular events include Nur1 phosphorylation (Figure 3), which accumulates upon DSB formation in rDNA repeats (Supplementary Figure 7). The effects of Nur1 phosphorylation are counteracted by the phosphatase Cdc14. It further requires Siz2-dependent SUMOylation of Nur1 and Lrs4, which earmarks the rDNA tethering complex for disassembly and occurs downstream of Nur1 phosphorylation (Figs. 5 and 6). Repeat release from the membrane-bound CLIP complex is ultimately mediated by the AAA-ATPase Cdc48/p97 in conjunction with its cofactor Ufd1 through recognition of SUMOylated Nur1 and Lrs4, which allows rDNA relocation and repair by Rad52 (Figs. 6 and 7).

The segregase Cdc48/p97 plays a key role in many cellular processes, including the DNA damage response, where it segregates ubiquitylated proteins from non-modified partner proteins to facilitate their proteasomal degradation. Recognition of ubiquitylated proteins is mediated by the heterodimeric cofactor Ufd1-Npl4, whereas other co-factors help Cdc48 to orchestrate the delivery to the proteasome (Franz et al., 2016). However, several studies revealed that Cdc48 also participates in disassembling protein complexes marked with SUMO (Bergink et al., 2013; Mérai et al., 2014; Nie et al., 2012). Consistent with these reports, we discovered that Cdc48 binds to SUMOylated Nur1 and Lrs4 through Ufd1 and disrupts the CLIP-cohibin complex to promote rDNA relocation upon damage (Figs. 6 and 7). Presenting multiple SUMOylated substrates may increase the efficiency of Cdc48^Ufd1-Npl4^-mediated CLIP-cohibin disassembly, and in agreement we found that overexpressing SUMO compensates for deficient rDNA recombination in the *nur1KR* mutant (Supplementary Figure 5). Additionally, we found that depletion of the p97 cofactor UFD1L impairs the nucleolar segregation of broken rDNA repeats in human cells (Figure 7). This implies that the role of Cdc48/p97^Ufd1-Npl4^ in rDNA release is conserved between yeast and humans, although the human substrate(s) remain(s) to be identified. Given the presence of mixed SUMO-ubiquitin chains, generated by STUbL E3 ligases, it has been proposed that Cdc48 can be recruited to its targets by a dual mechanism employing the SUMO- and ubiquitin-specific binding moieties of Ufd1 and Npl4, respectively (Franz et al., 2016; Nie et al., 2012). Besides recognizing SUMOylated proteins, Ufd1 has also been reported to bind directly to the fission yeast STUbL Rfp1 (homologous to *S. cerevisiae* Slx5/8 and human RNF4) (Køhler et al., 2013). We found that deletion of the STUbLs Slx5, Slx8 or Uls1 did not affect CLIP-cohibin dissociation (Supplementary Figure 6). However, we cannot exclude the possibility that STUbLs are involved downstream of CLIP-cohibin disassembly. It is thus noteworthy that the STUbL RNF4 acts together with UFD1L to release rDNA from the nucleolus upon damage (Figure 7), implying a functional link between ubiquitylation and p97 in relocating rDNA in human cells.

In addition to SUMO modification, Nur1 phosphorylation is another critical step controlling rDNA release. Specifically, we found that a mutant mimicking the phosphorylated state of Nur1 promotes DNA repair (Figure 3), whereas the corresponding mutant deficient in phosphorylation suppresses recombination (Figure 6). Nur1 is phosphorylated by the cyclin-dependent kinase Cdk1 to attenuate the release of the phosphatase Cdc14 during the cell cycle, which removes the modification in late anaphase (Godfrey et al., 2015; Holt et al., 2009). We found that Nur1 phosphorylation is also induced in response to DSBs (Supplementary Figure 7a). While CDK is inhibited after DNA damage in metazoans (Lanz et al., 2019), yeast Cdk1 was previously reported to be active at DSBs (Bantele et al., 2017; Huertas et al., 2008; Ira et al., 2004). It is therefore possible that this kinase is also involved in Nur1 phosphorylation during rDNA damage. Furthermore, cell cycle-dependent Nur1 phosphorylation might also serve as a signal to release the rDNA locus from the nuclear envelope to allow chromosome segregation, which would be counteracted by Cdc14 activity at the end of mitosis (Dauban et al., 2019; Godfrey et al., 2015). Alternatively, Nur1 could also be modified by DNA damage checkpoint kinases. For instance, ATM-mediated phosphorylation of Treacle (also known as *TCOF1*) is required to modulate the nucleolar response during rDNA damage in humans (Korsholm et al., 2019; Mooser et al., 2020). The molecular mechanism by which Nur1 phosphorylation triggers CLIP-cohibin disruption remains unknown. While SUMOylation does not affect Nur1 phosphorylation (Supplementary Figure 4), both appear to act in the same pathway (Figure 6), suggesting that Nur1 phosphorylation occurs upstream, and presumably controls SUMOylation. Consistent with this model, phosphorylation of other targets are known to stimulate their SUMOylation (Hietakangas et al., 2006; Kang et al., 2006).

Maintaining dynamic regulation of the CLIP-cohibin complex appears to be critical, since we find that tethering rDNA repeats permanently to the nuclear envelope is lethal (Figure 1). We propose that this is likely due to the failure to repair broken repeats, as the rDNA has been reported to be frequently damaged and relocated even in the absence of exogenous DNA damage (Torres-Rosell et al., 2007). However, we cannot exclude the possibility that permanent rDNA tethering also affects rDNA release during mitosis. Indeed, release of the *S. japonicus* CLIP complex from centromeres and telomeres is crucial for proper progression of mitosis (Pieper et al., 2020). Nonetheless, perinuclear anchoring of other genomic loci does not affect cell survival. For instance, artificial tethering of the mating-type loci in *S. cerevisiae* enhances transcriptional silencing without affecting cell growth (Andrulis et al., 1998). In agreement, we found that permanent perinuclear tethering of genomic regions recognized by the Gal4 transcription factor allowed normal growth (Figure 1). Together, these data suggest that maintaining a dynamic spatial regulation is specifically important for repetitive sequences, but independent of perinuclear release during mitosis in general.

In conclusion, we favor a model where phosphorylation and SUMOylation of the rDNA tethering complex act together to promote complex dissociation and modulate the release of broken repeats to be repaired (Figure 8). Remarkably, the spatial control of damaged rDNA repeats is highly reminiscent of other repair pathways, specifically the repair of DSBs within pericentromeric heterochromatin in flies and mice (Chiolo et al., 2011; Tsouroula et al., 2016). In this case HR progression is temporally blocked by SUMO and the Smc5/6 complex, whereas SUMOylation and ubiquitylation by STUbLs promotes the relocation of the damaged locus outside heterochromatin, and its targeting to the nuclear periphery for DNA damage repair (Chiolo et al., 2011; Ryu et al., 2015; Tsouroula et al., 2016). Our results provide another example of how cells utilize similar molecular mechanisms to control the relocation of repetitive sequences and thus maintain genome integrity.

**Figure 8.**
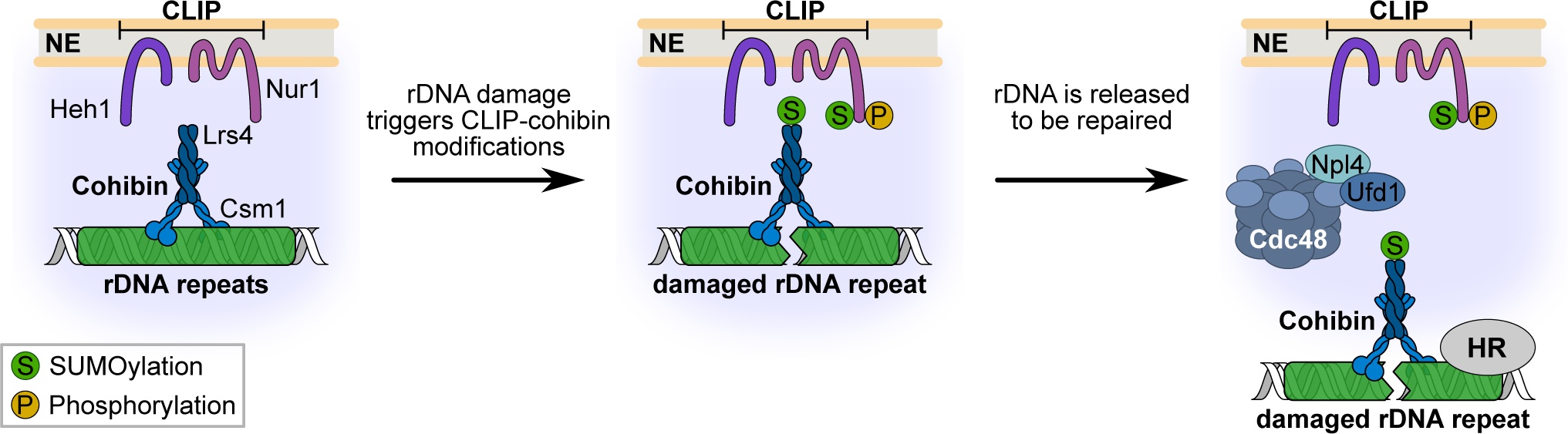
Model for rDNA release upon DNA damage. Under normal conditions, the rDNA repeats are kept inside the nucleolus by Cdc14-mediated dephosphorylation of Nur1, which maintains the interaction of the rDNA tethering complex. When a repeat unit is damaged, SUMOylation of CLIP-cohibin disrupts the complex through Cdc48-Npl4-Ufd1, allowing rDNA relocation outside the nucleolus. Nur1 is also phosphorylated, further supporting the broken repeat relocation.

### Methods Yeast strains

Yeast strains isogenic to W303 or DF5 were used for genetic studies, and PJ69-7a, Y187 or Y2H Gold for yeast two-hybrid assays. Full genotypes are listed in Supplementary Table S1.

### Yeast methods and molecular biology

Standard protocols for transformation, mating, sporulation and tetrad dissection were used for yeast manipulations (Sherman, 2002). Unless indicated otherwise, cells were grown at 30°C in YPD medium (1% yeast extract, 2% peptone and 2% glucose) or synthetic complete medium lacking individual amino acids to maintain selection for transformed plasmids. Protein tagging and the construction of deletion mutants were conducted by a PCR-based strategy and confirmed by immunoblotting and PCR, respectively (Janke et al., 2004; Knop et al., 1999). For galactose induction, cells were cultured in medium supplemented with 2% raffinose, and galactose was added to a mid-log phase yeast cells to a final concentration of 2%. Expression of Heh1 fusions, SUMO (Smt3) or Cdc48 was induced with galactose for 2h 30’. To generate DNA damage at the rDNA locus, I-*Sce*I and I-*Ppo*I expression was induced with galactose for 2h; for Lrs4 ChIP, I-*Ppo*I induction was performed for 1h 30’. For Nur1 C-terminal tagging with the active or inactive catalytic domain of SUMO-protease Ulp1 were performed using the pYM15-UD and pYM-uD^i^, respectively (Colomina et al., 2017). For auxin-inducible degradation, cells expressing the protein of interest fused to an auxin-dependent degron sequence derived from IAA17 together with the F-box protein TIR1 from *Oriza sativa* (Morawska and Ulrich, 2013) were cultured in YPD medium until mid-log phase, and the synthetic auxin analog 1-naphthaleneacetic acid (NAA) was added to a final concentration of 1.5 mM for 2h. Damage in rDNA repeats using I-*Ppo*I fused to the mouse glucocorticoid receptor was induced by adding galactose and triamcinolone acetonide (Wright et al., 1990) to a final concentration of 2% and 7.5 µM or 75 µM, respectively, to a mid-log phase yeast cells growing in minimal selective medium with 2% raffinose for 3h. Yeast growth assays were performed by spotting five-fold serial dilutions of the indicated strains on solid agar plates.

Cloning methods used standard protocols or the Gibson Assembly Master Mix (NEB). For yeast two-hybrid experiments, the different constructs were cloned into *pGADT7* or *pGBKT7* vectors (Clontech^TM^). Plasmids with point mutations or deletions were constructed by the PCR-based site-directed mutagenesis approach. Maps DNA sequences are available upon request.

### Yeast two-hybrid assays

The PJ69a strain was used to co-transform with the indicated plasmids, while the Y187 and Y2HGold strains were transformed with the indicated *pGBKT7* or *pGADT7* constructs, respectively, followed by mating. Spotting assays were performed on control medium (-LW) or selective medium with increased stringency (-LWH, -LWH + 1 mM 3-aminotriazol (3AT), -LWHA) and grown for 3 days, unless indicated otherwise.

### Live-cell imaging and analysis

For fluorescence microscopy, cells were diluted to OD_600_=0.15 in synthetic complete medium lacking individual amino acids, and grown until OD_600_=0.8-1.2 before imaging. Microscopy slides (MatTek) were pretreated with 1 mg ml^−1^ concanavalin A solution. Imaging was performed on a Zeiss AxioObserver Z1 confocal spinning-disk microscope equipped with an Evolve 512 (Photometrics) EMM-CCD camera through a Zeiss Alpha Plan/Apo 100×/1.46 oil DIC M27 objective lens. Optical section images were obtained at focus intervals of 0.2 µm. Subsequent processing and analyses of the images were performed in Fiji/ImageJ (Schindelin et al., 2012). For induction of I-*Sce*I endonuclease (rDNA damage), cells were grown at 30°C in synthetic complete medium lacking individual amino acids plus 2% raffinose to an OD_600_=0.6 and expression of I-*Sce*I was induced by addition of 2% galactose for 2h.

### Unequal sister chromatid exchange assays

Assays were performed as previously described (Huang et al., 2006), with some modifications. Cells were grown in YPDA or synthetic complete medium lacking individual amino acids to OD_600_=0.8-1.2, diluted 1:1,000 in distilled H_2_O and plated on synthetic complete medium plates. Cells were incubated at 30°C (for 3 days in glucose-containing medium, or for 4 days in galactose-containing medium) and then transferred to 4°C for 3 days to enhance color development. The rate of marker loss was calculated by dividing the number of half-red/half-white colonies by the total number of colonies. Red colonies were excluded from all calculations.

### Immunoblotting

Total protein extracts from 2−10^7^ cells (OD_600_=1) were prepared by trichloroacetic acid (TCA) precipitation (Knop et al., 1999). Proteins solubilized in HU loading buffer (8 M urea, 5% SDS, 200 mM Tris-HCl pH 6.8, 20 mM dithiothreitol (DTT) and bromophenol blue 1.5 mM) were resolved on NuPAGE 4%-12% gradient gels (Invitrogen), transferred onto polyvinylidene fluoride membranes (Immobilon-P) and analyzed by standard immunoblotting techniques using specific antibodies (see the ‘Antibodies’ section).

### Co-immunoprecipitation

Cell lysates were prepared by resuspending the pellets from 150-200 OD_600_ units in 800 µl lysis buffer (50 mM Tris pH 7.5, 150 mM NaCl, 10% glycerol, 1 mM EDTA pH 8, 0.5% NP-40, 1x complete EDTA-free protease inhibitor cocktail (Roche), 2 mM PMSF, 20 mM N-ethylmaleimide (NEM)). To block phosphatase activity, 1x tablet of PhosSTOP, 10 mM NaF and 20 mM ß-glycerophosphate were added to the lysis buffer. Cells were lysed by bead-beating (Precellys 24, Bertin instruments) with zirconia/silica beads (BioSpec Inc.) and lysates were cleared by centrifugation (800g, 5 min). Clarified extracts were incubated with pre-equilibrated antiHA-sepharose (Roche), GFP-Trap (Chromotek) or IgG-agarose beads (Sigma) for 1.5 hours at 4°C, beads were washed four times with lysis buffer and two times with wash buffer (50 mM Tris pH 7.5, 150 mM NaCl, 1 mM EDTA pH 8). Proteins were eluted by boiling with 30 µl HU loading buffer and analyzed by immunoblotting.

### Analysis of protein SUMOylation using semi-denaturing purification

To examine SUMOylated GFP-tagged proteins, cell lysates from 200 OD_600_ units from cells with or without rDNA damage were prepared as for co-immunoprecipitation experiments. Clarified extracts were incubated with 12.5 µl pre-equilibrated GFP-Trap beads (Chromotek) for 1.5 hours at 4°C, beads were washed once with lysis buffer. As reported for the detection of other post-translational modifications (Juretschke et al., 2019), the GFP-Trap beads were then washed five times with denaturation buffer (PBS, 8 M urea). The specifically bound proteins were eluted by boiling with 30 µl HU loading buffer and analyzed by immunoblotting.

### Analysis of protein SUMOylation by Ni-NTA pulldowns

To detect proteins covalently modified with SUMO, 200 OD_600_ units from cells expressing N-terminally histidine-tagged SUMO (6 histidines) under the control of the ADH1 promoter, integrated at the *URA3* locus, were collected. Subsequently, Ni-NTA pulldowns using Ni-NTA magnetic agarose beads (Qiagen) were performed as previously described (Psakhye and Jentsch, 2016).

### Generation of a SIM-binding deficient SUMO variant

To screen for SUMO interacting motif (SIM)-binding deficient mutants, we utilized strains overexpressing non-deconjugatable SUMO (SUMO^Q95P^) under control of the galactose inducible promoter *GAL1*, which decreased growth in WT cells. By contrast, SUMO^Q95P^ caused lethality in a strain lacking the SUMO-targeted ubiquitin ligase subunit Slx5 (*slx5Δ*). As Slx5 harbors multiple SIMs for substrate recognition, we assumed that mutating the SIM binding surface on SUMO^Q95P^ should cause lethality if expressed in WT cells, due to inhibited Slx5 interaction. In agreement to a previous report (Newman et al., 2017), a SIM-binding deficient mutation of the conserved F37 (Xie et al., 2010) in SUMO still allowed growth when expressed as a non-deconjugatable variant (SUMO^FA,Q95P^), suggesting that this SUMO variant still supports SUMO-SIM interaction. When we additionally mutated I39, WT cells expressing non-deconjugatable SUMO^FAIA,Q95P^ caused lethality, suggesting that Slx5 dependent clearance of this SUMO variant was blocked. The lost of SIM recognition was then assessed by co-immunoprecipitation of Slx5 with GFP-tagged SUMO^Q95P^ or the SUMO^Q95P^ variant F37A I39A.

### Chromatin immunoprecipitation (ChIP)

ChIP experiments were performed as described previously (Braun et al., 2011), with some modifications. Briefly, 150 ml cell cultures were grown to OD_600_=0.8-1.0, cross-linked (1% formaldehyde, 16 min, room temperature) and lysed by bead beating. The chromatin fraction was isolated and sheared to 200bp - 500bp fragments (30 min, 30-sec on/off cycles, 90% amplitude) using a Q800R1 sonicator (QSonica). Immunoprecipitations were performed overnight at 4°C with 100 µl of 25% slurry of prewashed IgG-agarose beads (Sigma) from lysates corresponding to 80-100 OD_600_ of cells. The beads were washed and eluted, and the eluate was reverse cross-linked at 65°C for 3 h and incubated with proteinase K for 2 h at 55°C. DNA was cleaned up with ChIP DNA Clean & Concentrator^TM^ kit (#D5201, Zymo Research). Immunoprecipitated DNA was quantified by qPCR using primaQUANT SYBR Master mix (Steinbrenner Laborsysteme GmbH. #SL-9902B) and a QuantStudio^TM^ 3/5 Real-Time PCR system (Applied Biosystems/Thermo Fisher). Primers are listed in Supplementary Table S2. To avoid changes due to different rDNA copy number, the relative fold enrichment for each strain was determined over the average of three rDNA positions (RDN1 #9, RDN1 #12 and RDN1 #25), after normalization to input: [rDNA(IP)/rDNA(WCE)]/[mean(rDNA)(IP)/mean(rDNA)(WCE)].

### Determination of rDNA copy number and induction of double-strand breaks

The amount of cells corresponding to OD_600_=1 (2−10^7^ cells) was harvested before and after induction of rDNA damage using I-*Ppo*I (uninduced and induced samples, respectively), and genomic DNA was prepared using the MasterPure Yeast DNA Purification Kit (Epicenter). Genomic DNA was then used as input for qPCR with specific primers (Supplementary Table S2). The relative rDNA copy number or intact I-*Ppo*I cut site (rDNA damage efficiency) was calculated as the ratio between induced and uninduced samples, after normalized to a control locus on chromosome VI (*ACT1*).

### Quantitative RT-qPCR

RT-qPCR analyses were performed as previously described (Braun et al., 2011). cDNAs were quantified by qPCR using primaQUANT SYBR Master mix (Steinbrenner Laborsysteme GmbH) and a QuantStudioTM 3 Real-Time PCR system (Applied Biosystems/Thermo Fisher). Primers are listed in Table S1. For the calculation of mean values and standard deviation from independent experiments, *ACT1*-normalized data sets are shown in log_2_ scale as relative to the mean value of the wild type.

### Cell culture

RPE cells were grown in DMEM (Sigma) supplemented with Penicillin-Streptomycin (Life Technologies), L-Glutamine (ThermoFisher) and 10% fetal bovine serum (FBS) (Takarabio) at 37°C and 5% CO_2_. RNAi experiments were performed using RNAiMax (Life Technologies) two days before treatments. The following siRNA sequences were used: siControl (5’-UGGUUUACAUGUCGACUAA-3’; Metabion), siUFD1La (5’-GUGGCCACCUACUCCAAAUUU-3’; Metabion), siUFD1Lb (5’-CUACAAAGAACCCGAAAGAUUUU-3’; Metabion). ^HA^I-*Ppo*I expression was induced by adding doxycycline (1 µg/ml, Sigma) 4h and Shield-1 (1 µM, Aobious) 1h before fixation.

### Immunofluorescence and quantification of nucleolar release in human cells

Cells were washed with PBS (Sigma) and fixed in 4% paraformaldehyde for 30 min at room temperature. Then, cells were permeabilized in CSK buffer (10 mM HEPES pH 7.4, 300 mM Sucrose, 100 mM NaCl, 3 mM MgCl_2_) with 0.1% Triton-X for 30 min. Blocking was performed for 1 h with 5% FBS in PBS after which cells were stained in 1% FBS in PBS overnight at 4°C. After primary staining, cells were washed 3 times 5 min with PBS and incubated with secondary antibodies for 1h at RT. After another three 5 min washes in PBS, slides were mounted with Aqua-Poly/Mount (Polysciences). Images were obtained using an AxioObserver Z1 confocal spinning-disk microscope (Zeiss) equipped with an AxioCam HRm CCD camera (Zeiss) or a sCMOS ORCA Flash 4.0 camera (Hamamatsu) and a Plan/Apo 63 Å∼/1.4 water-immersion objective. The percentage of cells with nucleolar retention of γH2AX foci were defined as cells that show an overlap (yellow) of the γH2AX (green) with the NPM (red) signal. Blinded visual scoring of rDNA damage retention was performed by two individuals.

### Antibodies

Polyclonal Smt3 (1:5,000), Slx5 (1:5,000) and Cdc48 (1:5,000) antibodies were raised in rabbits and have been described previously (Hoege et al., 2002; Höpfler et al., 2019; Richly et al., 2005). Monoclonal antibodies directed against the HA epitope (1:1,000; 3F10) and monoclonal antibody against GFP (1:1,000; B-2) were purchased from Roche and Santa Cruz Biotechnology, respectively. Mouse monoclonal antibodies against Dpm1 (1:2,000; 5C5A7) and Pgk1 (1:5,000; 22C5D8) were obtained from Invitrogen. Mouse monoclonal antibodies against phospho-H2AX (Ser139) (1:1,000, JBW301) were obtained from Merck Millipore. Rabbit polyclonal antibodies against Nucleophosmin (1:100, ab15440) and Rad53 (1:1000, ab104232) were purchased from Abcam. Secondary antibodies goat-α-mouse IgG Alexa fluor 488 (1:5,000; A11001) and donkey-α-rabbit IgG Alexa fluor 568 (1:5,000; A10042) were obtained from Thermo Fisher Scientific.

### Statistics and reproducibility

Representative results of at least two independent experiments were presented in all of the figure panels. Analyses of the variance (ANOVA) were performed, and pairwise differences were evaluated with Tukey’s *post hoc* test using R statistical language (R Development Core Team, 2008); different groups are marked with letters at the 0.05 significance level. For all error bars, data are mean ± S.E.M. P values for all graphs were generated using two-tailed Student’s t-tests; N/S, P ≥ 0.05, *P ≤ 0.05, **P ≤ 0.01, ***P ≤ 0.001.

## Acknowledgements

We thank members of the Braun and Ladurner labs for fruitful discussions; S. Lall from LSE editors for critical comments on the manuscript; R. H. Medema for RPE cell lines; J. Torres-Rosell for *pYM15-UD* and *pYM15-uD^i^* plasmids; P. Bourilhon and D. Sinitski for mouse cDNA and reagents; S. Fischer-Burkart, T. S. van Emden and A. Muhammad for assistance, especially during COVID-19 times. This study was supported by a grant awarded to S.B. from the German Research Foundation (DFG; BR 3511/4-1). I.K.M. is recipient of a grant from the Stiftungskommission of the LMU Medical Faculty. Research in the S.J. laboratory was supported by Max Planck Society (MPG), Center for Integrated Protein Science Munich (CIPSM), Louis-Jeantet Foundation, and a European Research Council (ERC) Advanced Grant. Research in the S.B., B.P., and A.G.L. laboratories was further supported by the DFG through the collaborative research center CRC 1064 (project ID 213249687-SFB 1064) and by the LMU (A.G.L.) and MPG (B.P.).

## Author contributions

M.C., S.J., and S.B. conceived the study and designed experiments. I.K.M. performed the mammalian experiments. L.M.C. contributed to the microscopy studies in yeast. F.d.B. designed the SUMO F37A I39A mutant. M.C. performed all other experiments. M.C., I.K.M., and L.M.C. analyzed the data. S.B., and S.J. supervised the project. S.B., S.J., B.P., and A.G.L. acquired funding. M.C., and S.B. conceived and wrote the manuscript, and all authors contributed to editing.

## Supplementary information

**Supplementary Figure 1.**
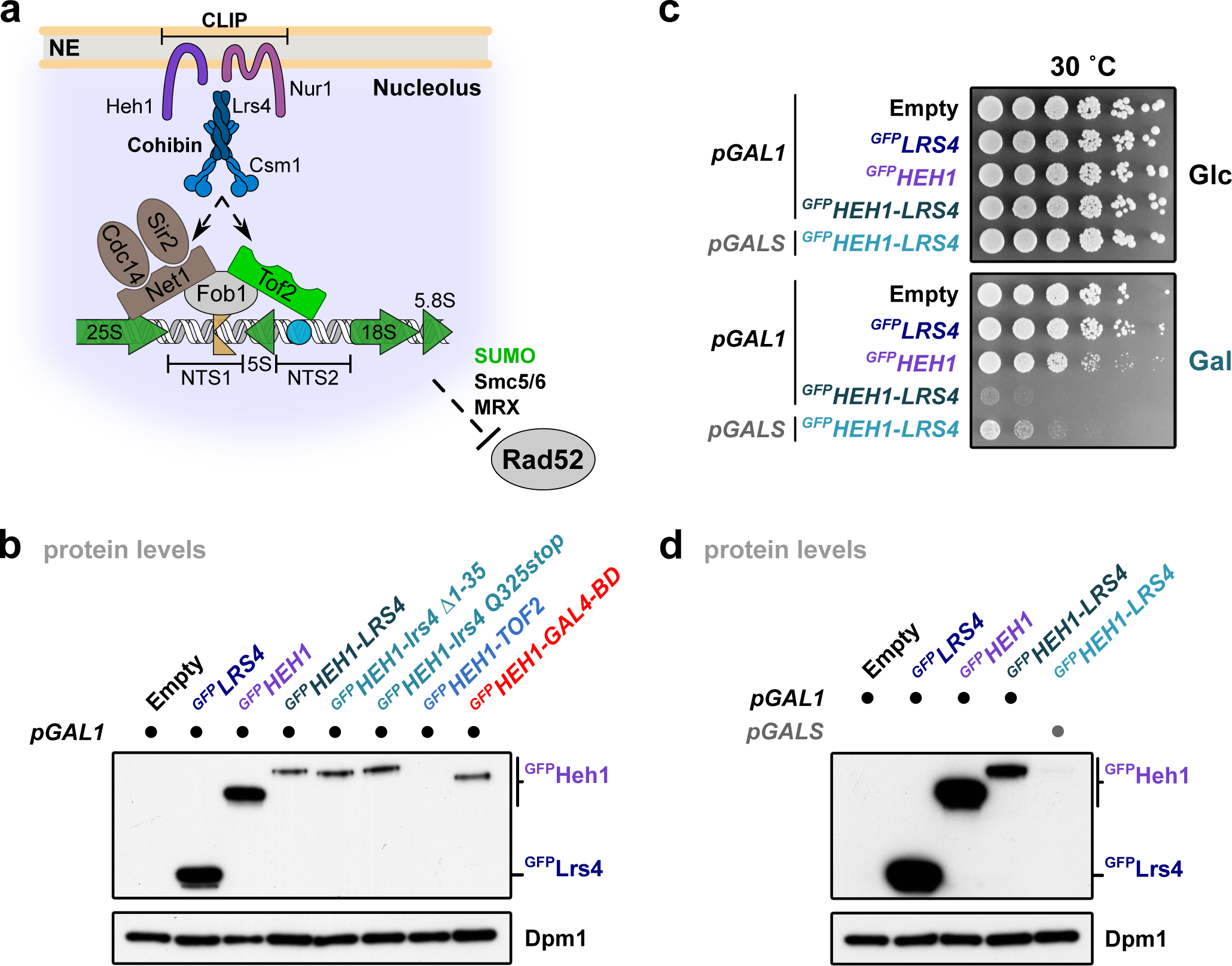
Disruption of rDNA dynamics causes severe growth defects in yeast. **a**, Scheme of an rDNA unit bound by the tethering complex in *Saccharomyces cerevisiae*. Each repeat of 9.1 kb presents a near-identical sequence comprising the 35S (subdivided in 5.8S, 18S and 25S) and 5S ribosomal RNA coding genes. Polymerase II silenced regions are known as NTS1 (non-transcribed spacer 1) and NTS2 regions, which are located down- and upstream of the 5S gene, respectively. NTS1 contains the replication fork barrier (RFB), whereas NTS2 contains an autonomous replicating sequence (ARS). The rDNA units are attached to the inner nuclear membrane via interaction between cohibin (Lrs4 and Csm1) and CLIP (Heh1 and Nur1). Cohibin then binds to the NTS1 region of the rDNA repeats through an rDNA-bound protein complex that includes Fob1, Tof2 and the RENT complex (regulator of nucleolar silencing and telophase exit; composed of Cdc14, Net1 and Sir2). SUMO together with the MRX and Smc5/6 complexes exclude Rad52 from the nucleolus. **b**, Immunoblot of WT cells transformed with empty vector or plasmids bearing GFP-tagged *LRS4*, *HEH1* or fusion proteins with the galactose-inducible promoter after induction. **c**, Five-fold serial dilutions of WT cells transformed with empty vector or plasmids bearing the indicated GFP-tagged fusion proteins with different galactose-inducible promoters (either *pGAL1* or *pGALS*), as indicated. Cells were spotted and grown on selective media with glucose (control, Glc) or galactose (induction, Gal) at 30°C for 3 days. **d**, Immunoblot of the strains used in **c**, after galactose induction. For **b** and **c**, the different strains were grown to mid-log phase, fusion proteins were induced by adding 2% galactose for 2 h 30’. Dpm1 served as loading control.

**Supplementary Figure 2.**
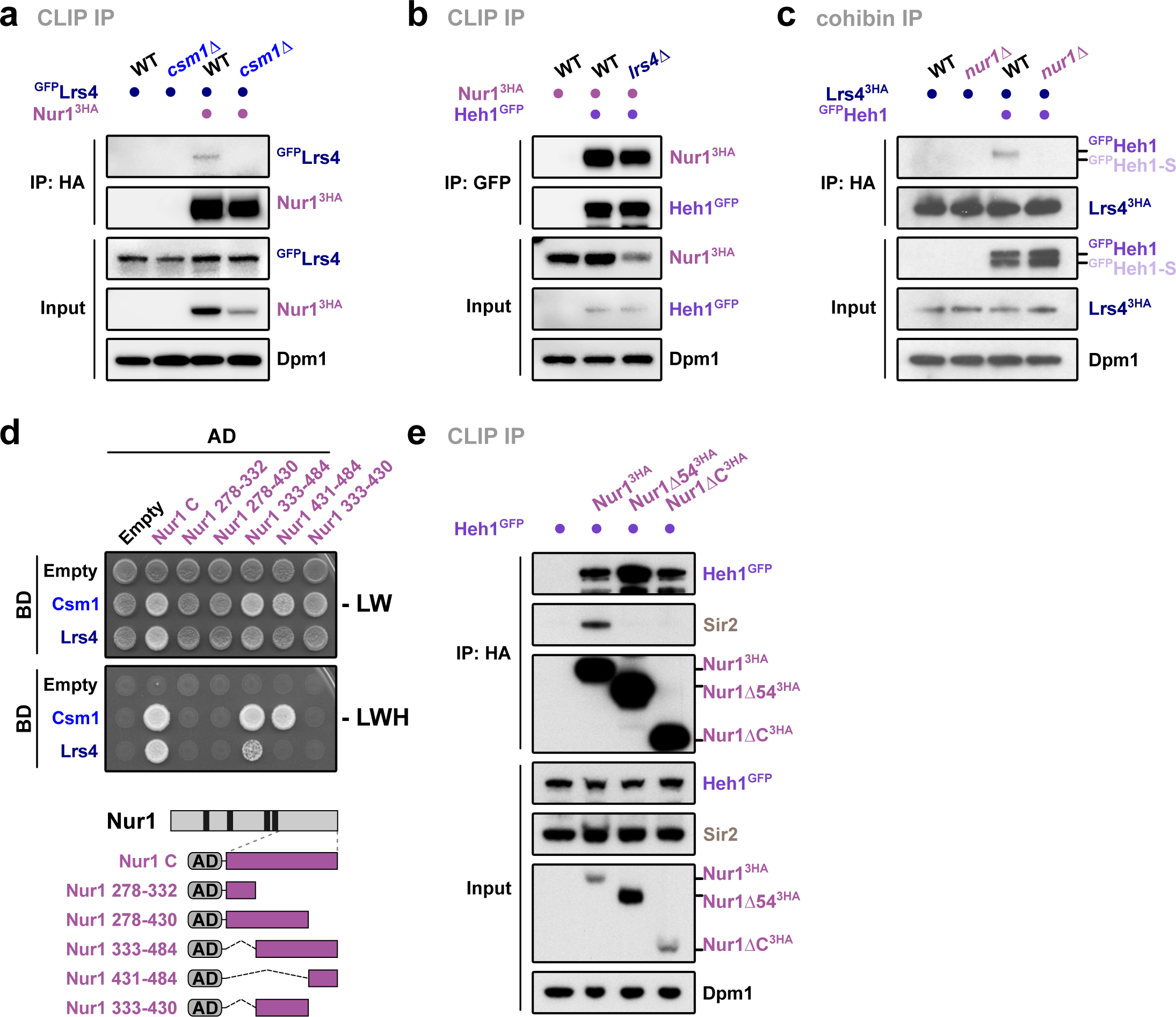
The C-terminal residues of Nur1 are required for CLIP-cohibin complex formation. **a** to **c**, Co-immunoprecipitation of ^GFP^Lrs4 with Nur1^3HA^ in WT or *csm1Δ* cells (**a**); of Nur1^3HA^ with Heh1^GFP^ in WT or *lrs4Δ* cells (**b**); and of ^GFP^Heh1 with Lrs4^3HA^ in WT and *nur1Δ* cells (**c**). Due to N-terminal tagging of Heh1, both spliced versions (denoted as Heh1 and Heh1-S) can be detected in **c**. **d**, Y2H analysis of Csm1 and Lrs4 with either Nur1 C or truncated mutants. Fusions with Gal4-activating domain (AD) or Gal4-DNA-binding domain (BD) are indicated. Schematic showing Nur1 truncated constructs. **e**, Co-immunoprecipitation of Heh1^GFP^ and Sir2 with either HA-tagged Nur1 full-length (Nur1^3HA^), lacking its last 54 residues (Nur1Δ54^3HA^) or the complete C-terminal domain (Nur1ΔC^3HA^). For immunoblots, Dpm1 served as loading control. For Y2H assays, cells were spotted on control media (-LW) or selective media (-LWH) and grown for 3 days.

**Supplementary Figure 3.**
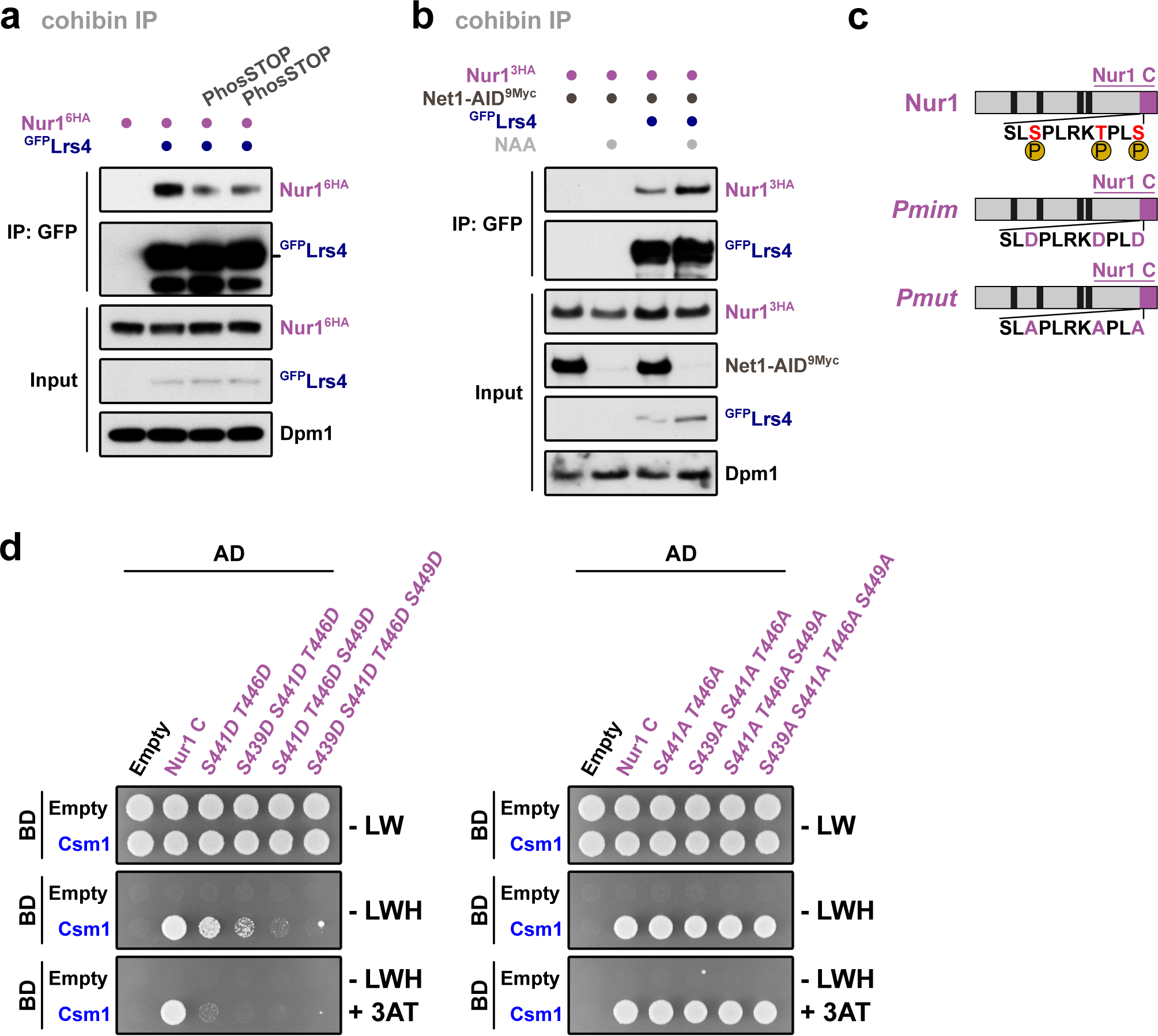
Phosphorylation of C-terminal Nur1 impairs CLIP-cohibin interaction. **a**, Co-immunoprecipitation of Nur1^3HA^ with ^GFP^Lrs4 in WT with impaired global phosphatase activity (PhosSTOP). One untreated and two independently treated samples are shown. **b**, Co-immunoprecipitation of Nur1^3HA^ with ^GFP^Lrs4 in WT or Net1-depleted cells. Degradation of a C-terminal AID (auxin inducible degron) fusion of Net1 (Net1-AID^9Myc^) was induced through treatment with 1-naphthaleneacetic acid 1.5 mM (NAA, a synthetic analog of auxin) for 2 h. **c**, Schematic highlighting Nur1 residues important for CLIP-cohibin interaction in red. Amino acids 439 to 449 are depicted. P, phosphorylation. **d**, Y2H analysis of Csm1 (BD) with Nur1 C and phosphomimetic (left) or phosphomutant (right) constructs (AD), as indicated. For immunoblots, Dpm1 served as loading control. For Y2H assays, cells were spotted on control media (-LW) or selective media (-LWH or -LWH + 3AT 1 mM) and grown for 3 days.

**Supplementary Figure 4.**
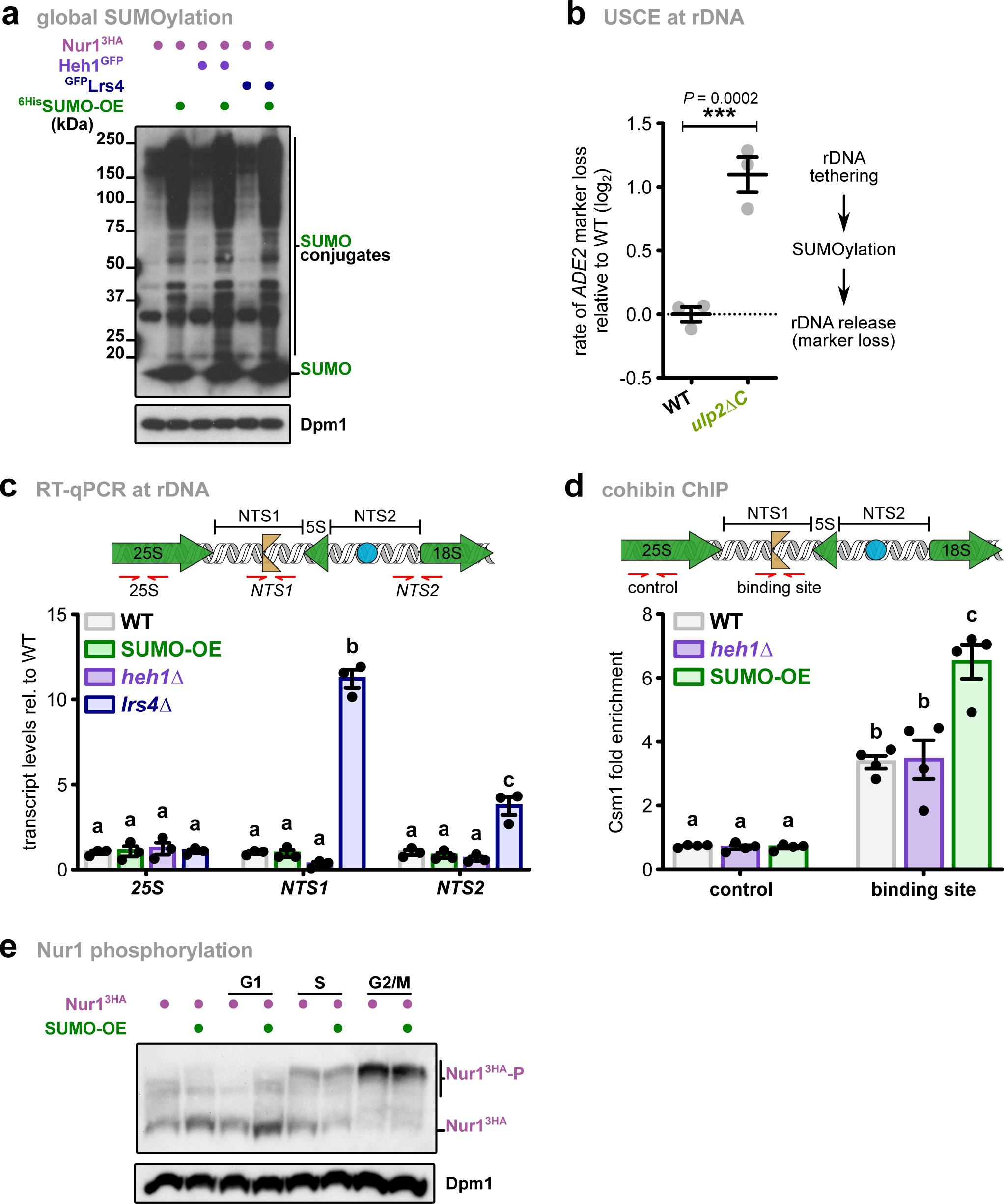
Overexpression of SUMO increases rDNA instability by releasing the repeats from the nucleolus. **a**, Immunoblot of SUMO conjugates of cells from Figure 4A, expressing SUMO at endogenous levels or overexpressed from the *ADH1* promoter (^6His^SUMO-OE). **b**, Rates of unequal rDNA recombination in WT and *ulp2ΔC* cells. The rate of marker loss is calculated as the ratio of half-sectored colonies to the total number of colonies, excluding completely red colonies; data is shown in log_2_ scale relative to WT. **c**, RT-qPCR analysis of WT, SUMO-OE, *heh1Δ* and *lrs4Δ* strains. Shown are transcript levels relative to WT after normalization to *ACT1*. Illustration shows a schematic representation of an rDNA unit. Amplified regions are highlighted with red arrows. NTS, non-transcribed spacer; yellow symbol, replication fork block; blue symbol, ARS. **d**, Csm1TAP binding to rDNA repeats in WT, *heh1Δ* and SUMO overexpressing (SUMO-OE) cells, quantified by ChIP-qPCR. The ChIP values are shown as Csm1^TAP^ fold enrichment over the average of three rDNA positions, after normalization to input. Scheme of an rDNA repeat as in **c** is shown; in red are marked the amplified regions. **e**, Nur1^3HA^ phosphorylation-dependent mobility shifts in cells expressing SUMO at endogenous levels or overexpressed from the *ADH1* promoter (^6His^SUMO-OE), analyzed using Phos-tag gels. Cells were arrested in G1, S and G2/M by treating the cells with α-factor, HU 100 mM or nocodazole 5 ug/ml for 2h, respectively. For immunoblots, Dpm1 served as loading control. For **b**, **c** and **d**, data are mean of n = 3-4 independent biological replicates. Analysis of variance (ANOVA) was performed, and different letters denote significant differences with a Tukey’s *post hoc* test at P < 0.05.

**Supplementary Figure 5.**
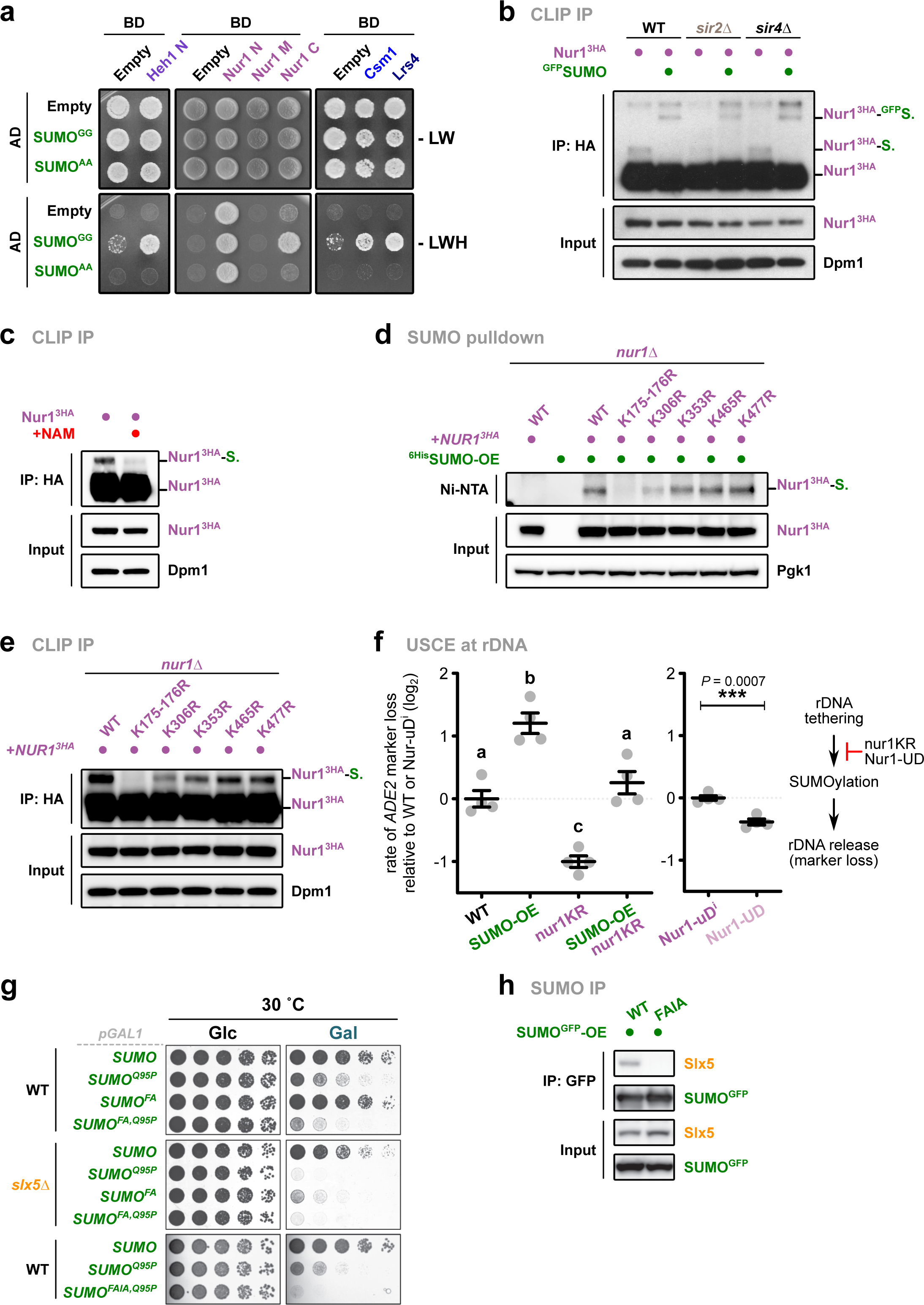
SUMOylation of CLIP-cohibin members affects rDNA stability. **a**, Y2H analysis of (SUMO^GG^) or a conjugation-deficient SUMO (SUMO^AA^) with Heh1 N and Nur1 nucleoplasmic domains, Csm1 and Lrs4. Fusions with Gal4-activating domain (AD) or Gal4-DNA-binding domain (BD) are indicated. Cells were spotted on control media (-LW) or selective media (-LWH) and grown for 3 days. **b**, Immunoprecipitation of Nur1^3HA^ in WT, *sir2Δ* and *sir4Δ* expressing SUMO (endogenous promoter) or ^GFP^SUMO (*ADH1* promoter). Bands corresponding to Nur1 unmodified or monoSUMOylated are labeled. **c**, Immunoprecipitation of Nur1^3HA^ in WT cells treated with nicotinamide (NAM) 5 mM for 2h. Bands corresponding to Nur1 unmodified or monoSUMOylated are labeled. **d**, Immunoprecipitation of Nur1^3HA^ in *nur1Δ* cells transformed with plasmids bearing *NUR1* or the indicated mutants with the endogenous *NUR1* promoter. Bands corresponding to Nur1 unmodified or monoSUMOylated are labeled. **e**, Denaturing Ni-NTA pulldowns of ^6His^SUMO conjugates from *nur1Δ* cells transformed with plasmids bearing *NUR1* or the indicated mutants with the endogenous *NUR1* promoter. Pgk1 served as loading control. **f**, Rates of unequal rDNA recombination in WT cells expressing Nur1 or the SUMOylation deficient mutant *nur1KR* with endogenous or increased levels of SUMO, grown in galactose-containing media (left); and of strains expressing Nur1 fused to the SUMO-protease domain of Ulp1 (Nur1-UD) or the inactive version (Nur1-uD^i^), which carries the F474A and C580S mutations (right). For SUMO overexpression, the different strains were transformed with empty vector or a plasmid bearing SUMO with the GAL1 promoter. The rate of marker loss is calculated as the ratio of half-sectored colonies to the total number of colonies, excluding completely red colonies, shown in log_2_ scale relative to WT (left) or Nur1-uD^i^ (right); data are mean of n = 4 independent biological replicates. Analysis of variance (ANOVA) was performed, and different letters denote significant differences with a Tukey’s post hoc test at P < 0.05. g, Five-fold serial dilutions of WT or slx5Δ cells transformed with plasmids bearing SUMO WT and the indicated SUMO variants with the galactose-inducible promoter *GAL1*. Cells were spotted and grown on selective media with glucose (control, Glc) or galactose (induction, Gal) at 30°C for 3 days. h, Co-immunoprecipitation of Slx5 with either GFP-tagged SUMO WT or a mutant unable to recognize SIMs (FAIA, which harbors the mutations F37A I39A). For **b**, **c** and **e**, Dpm1 served as loading control.

**Supplementary Figure 6.**
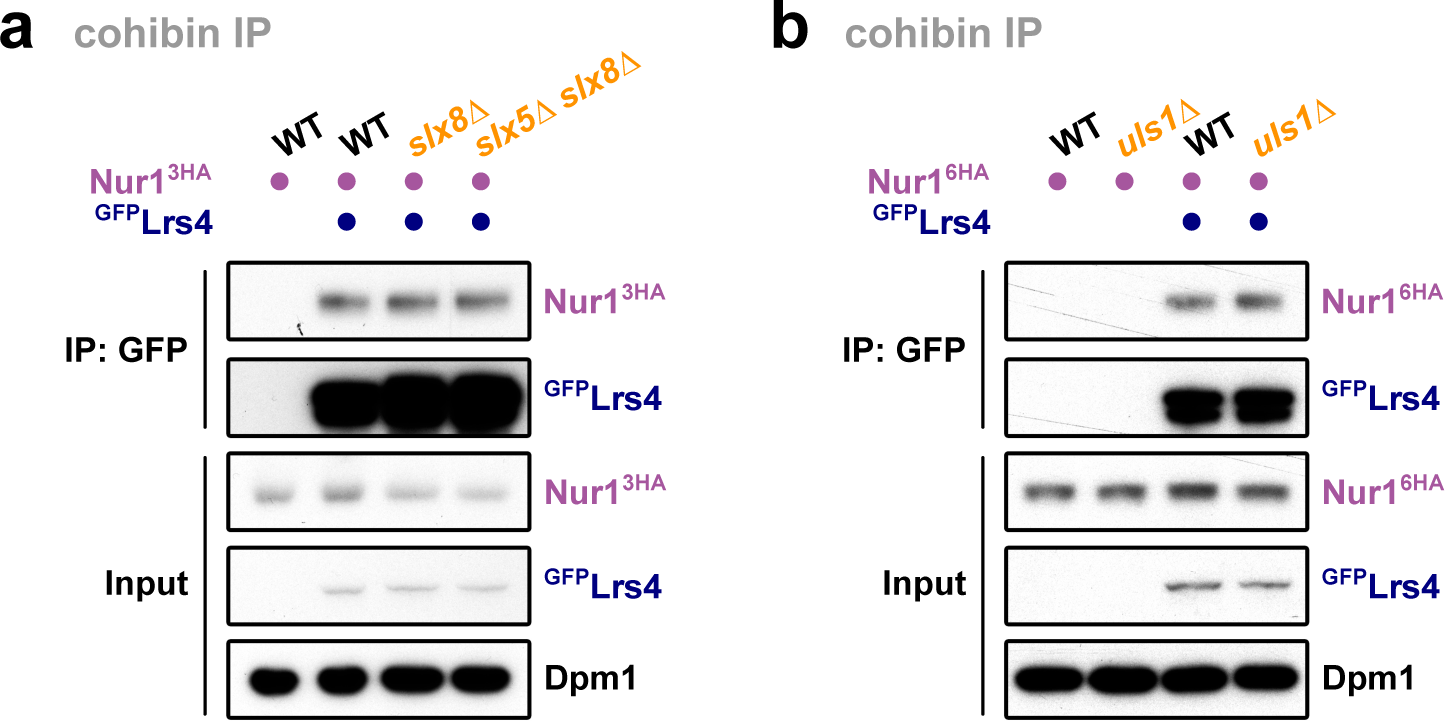
CLIP-cohibin interaction is not affected by the STUbLs. **a** and **b**, Co-immunoprecipitation of HA-tagged Nur1 with ^GFP^Lrs4 in WT, *slx8Δ*, *slx5Δ slx8Δ* (**a**), or *uls1Δ* (**b**) cells, as indicated. Dpm1 served as loading control.

**Supplementary Figure 7.**
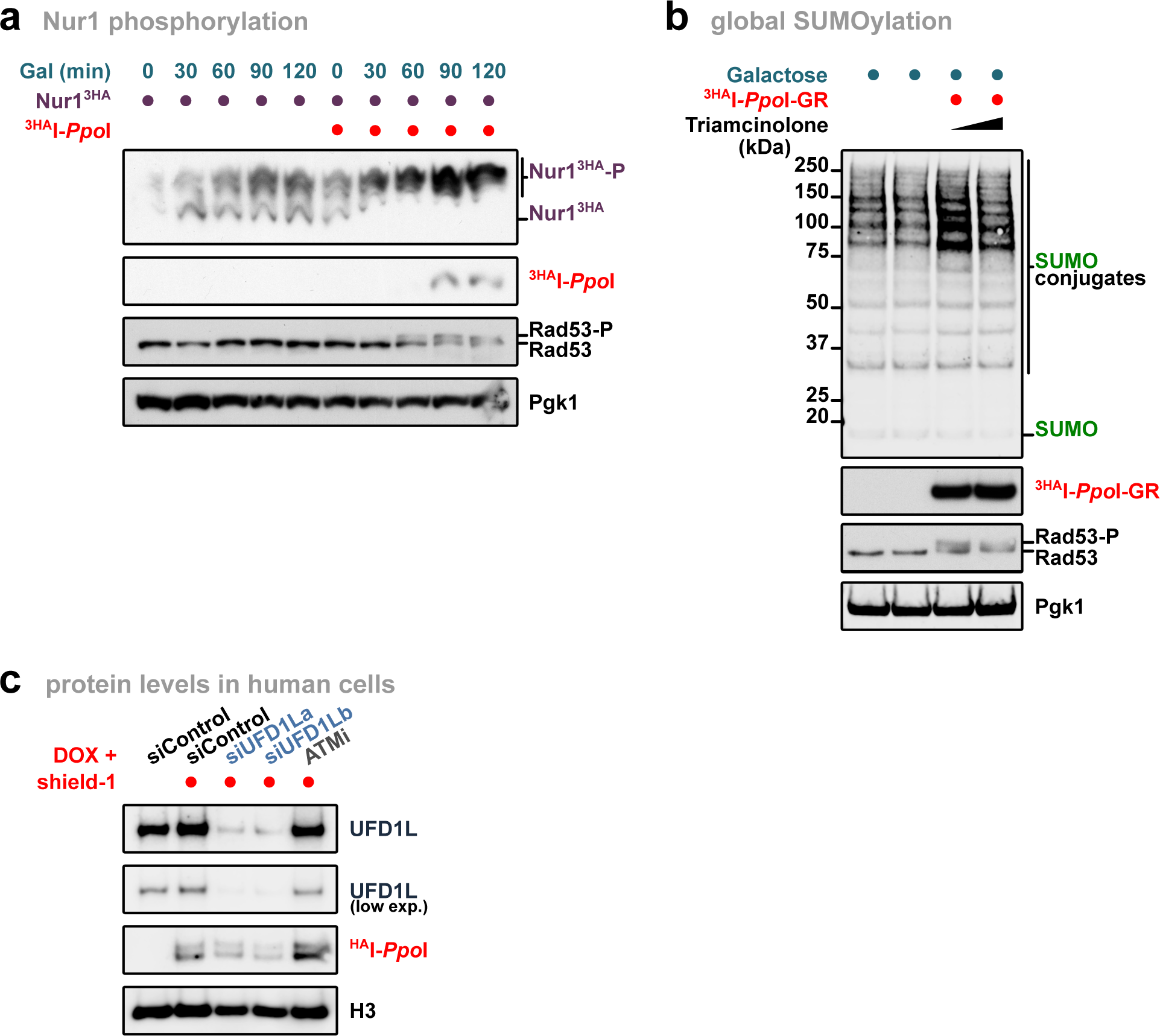
Damage at rDNA promotes release of broken repeats. **a**, Nur1^3HA^ phosphorylation-dependent mobility shifts cells in cells with or without rDNA damage, analyzed using Phos-tag gels. Dpm1 served as loading control. ^3HA^I-*Ppo*I with the galactose-inducible promoters *GALL* were integrated at the *LEU2* locus. The different strains were grown to mid-log phase, and the endonuclease was induced by adding 2% galactose for indicated times. **b**, Immunoblot of SUMO conjugates upon rDNA damage. To ensure tight regulation of the endonuclease, I-*Ppo*I was fused to the glucocorticoid receptor ligand-binding domain (^3HA^I-*Ppo*I-GR) and expressed under the galactose-inducible promoter GAL1 from a centromeric plasmid *YCplac22*. The strains were grown to mid-log phase, and the endonuclease was induced by adding 2% galactose for 3 h. Nuclear location of ^3HA^I-*Ppo*I-GR was induced by addition of triamcinolone acetonide 7.5 μM and 75 μM. **c**, Immunoblot of RPE cells from Fig. 7d after rDNA damage, showing knock-down efficiency of UFD1L and ^HA^I-*Ppo*I induction. H3 served as loading control. For **a** and **b**, Rad53 phosphorylation served as a marker for DNA damage, while Pgk1 served as loading control.

**Supplementary Table S1.**
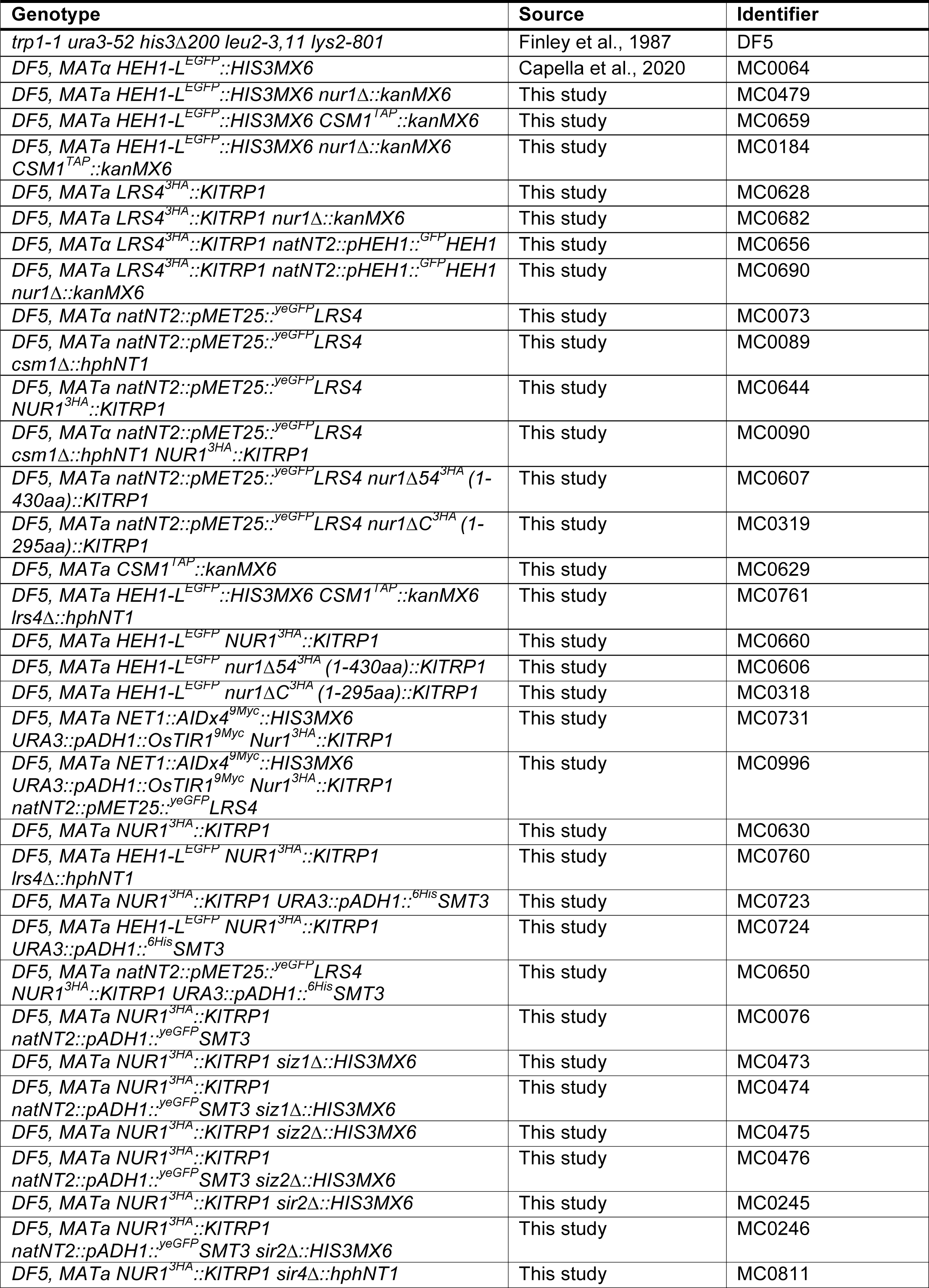
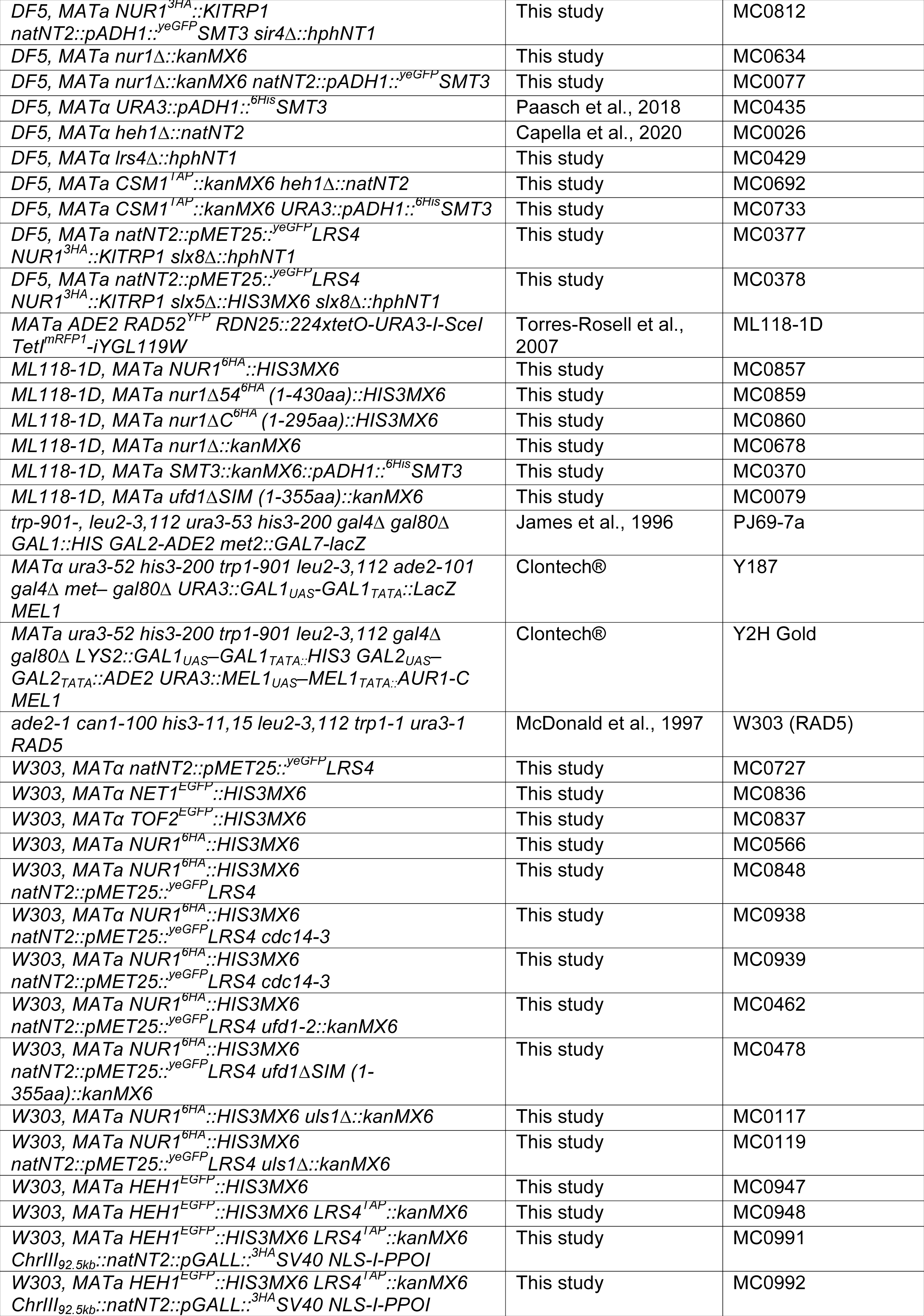
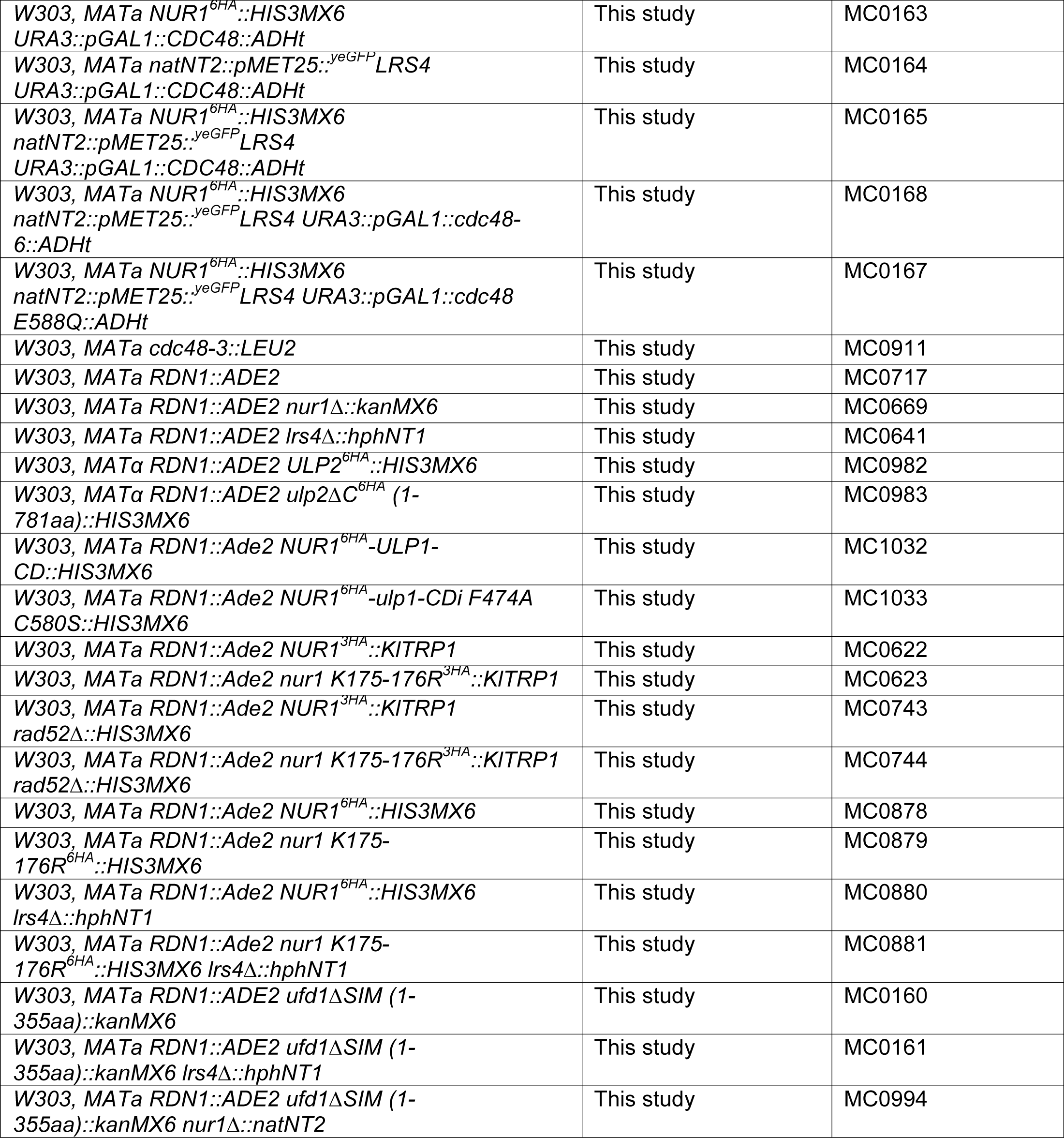
Yeast strains used in this study.

---

**Supplementary Table S2.**
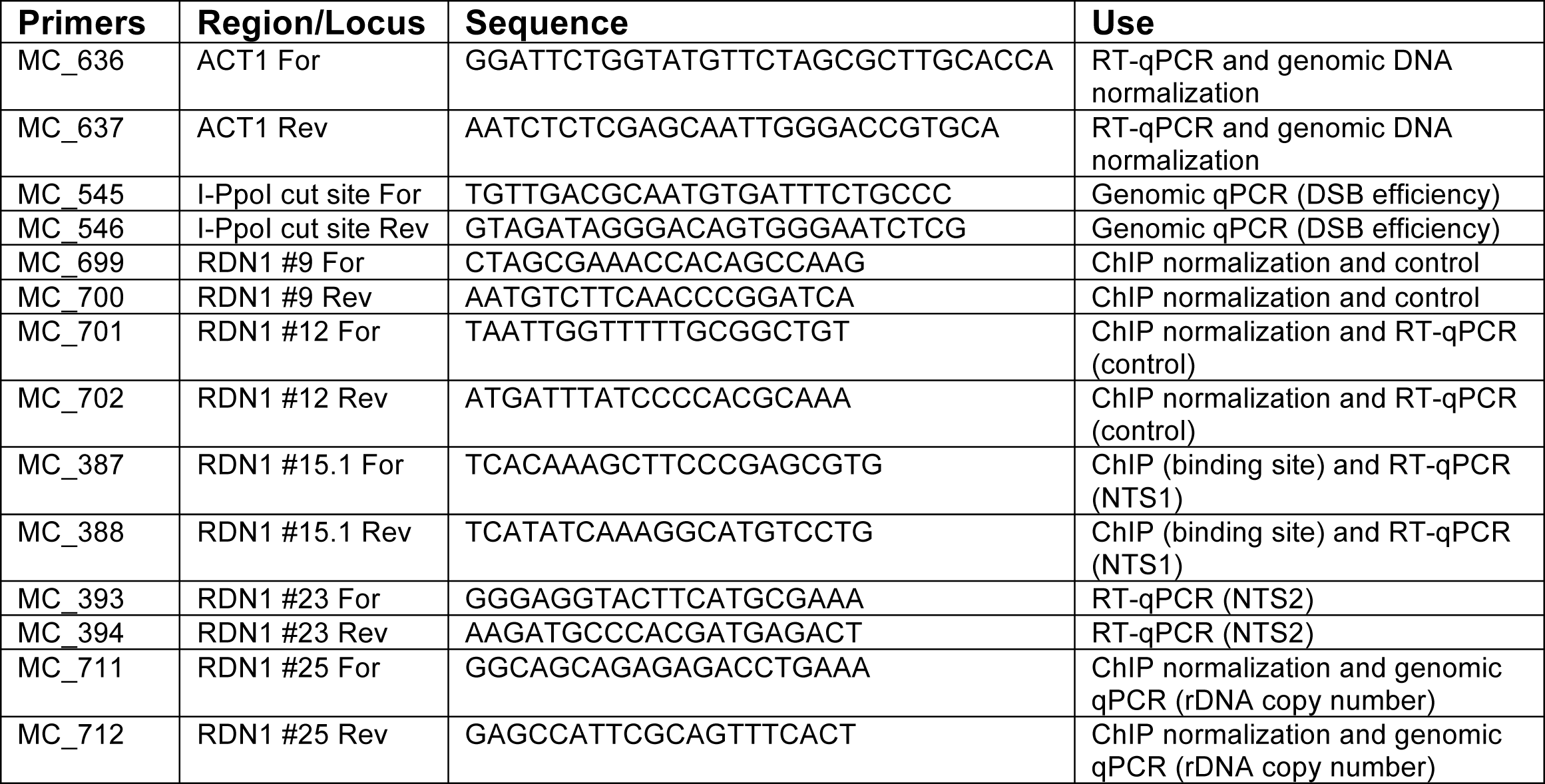
Set of primers used in this study for qPCR or ChIP experiments.

**Supplementary Table S3.** Plasmids used in this study. Point mutations refer to the position of the aminoacid/s (aa) in WT proteins. The specific positions of the truncated constructs are indicated in brackets.

| Construct | Source | Identifier |
| --- | --- | --- |
| YCplac22 | Gietz and Akio, 1988 | pMC_0111 |
| YCplac22 pGAL1::HEH1-L-3HA-GBP::ADH1t | This study | pMC_0442 |
| YCplac22 pNOP1::NOP1-ECFP | This study | pMC_0480 |
| YCplac22 pGAL1::NUR1::ADH1t | This study | pMC_0595 |
| YCplac22 pGAL1::nur1 S441D T446D S449D::ADH1t | This study | pMC_0599 |
| YCplac22 pGAL1::nur1 S441A T446A S449A::ADH1t | This study | pMC_0600 |
| YCplac111 | Gietz and Akio, 1988 | pMC_0110 |
| YCplac111 pGAL1::yeGFP-HEH1-L::ADH1t | Capella et al., 2020 | pMC_0395 |
| YCplac111 pGAL1::yeGFP-LRS4::ADH1t | This study | pMC_0631 |
| YCplac111 pGAL1::yeGFP-HEH1-L-LRS4::ADH1t | This study | pMC_0443 |
| YCplac111 pGAL1::yeGFP-HEH1-L-lrs4 Δ1-35aa::ADH1t | This study | pMC_0444 |
| YCplac111 pGAL1::yeGFP-HEH1-L-lrs4 Q325Stop::ADH1t | This study | pMC_0445 |
| YCplac111 pGAL1::yeGFP-HEH1-L-TOF2::ADH1t | This study | pMC_0446 |
| YCplac111 pGAL1::yeGFP-HEH1-L-GAL4-BD::ADH1t | This study | pMC_0447 |
| YCplac111 pGALS::yeGFP-HEH1-L-LRS4::ADH1t | This study | pMC_0448 |
| YCplac111 pNUR1::NUR1::ADH1t | This study | pMC_0565 |
| YCplac111 pNUR1::nur1 S441D T446D S449D::ADH1t | This study | pMC_0570 |
| YCplac111 pNUR1::NUR1-TAP::ADH1t | This study | pMC_0565 |
| YCplac111 pNUR1::nur1 S441D T446D S449D-TAP::ADH1t | This study | pMC_0570 |
| YCplac111 pNUR1::NUR1-3HA | This study | pMC_0545 |
| YCplac111 pNUR1::nur1 K175-176R-3HA | This study | pMC_0546 |
| YCplac111 pNUR1::nur1 K306R-3HA | This study | pMC_0547 |
| YCplac111 pNUR1::nur1 K353R-3HA | This study | pMC_0548 |
| YCplac111 pNUR1::nur1 K465R-3HA | This study | pMC_0549 |
| YCplac111 pNUR1::nur1 K477R-3HA | This study | pMC_0550 |
| YCplac111 pNUR1::smt3 Q95P-NUR1-3HA | This study | pMC_0555 |
| YCplac111 pNUR1::smt3 F37A I39A Q95P-NUR1-3HA | This study | pMC_0556 |
| YCplac111 pLRS4::LRS4-TAP::ADH1t | This study | pMC_0637 |
| YCplac111 pLRS4::smt3 Q95P-LRS4-TAP::ADH1t | This study | pMC_0638 |
| YCplac111 pLRS4::smt3 F37A I39A Q95P-LRS4-TAP::ADH1t | This study | pMC_0639 |
| YCplac111 pNUR1::NUR1-3HA | This study | pMC_0545 |
| YCplac111 pNUR1::smt3 Q95P-NUR1-3HA | This study | pMC_0555 |
| YCplac111 pNUR1::smt3 F37A I39A Q95P-NUR1-3HA | This study | pMC_0546 |
| YCplac111 pGAL1::2xSV40 NLS-HA-I-SceI::ADH1t | This study | pMC_0298 |
| pRS415 pGAL1 | Mumberg et al., 1995 | pMC_0147 |
| pRS415 pGAL1::3HA-I-Ppol-GR-LBD::CYC1t | This study | pMC_0320 |
| pRS415 pTEF1 | This study | pMC_0485 |
| pRS415 pTEF1::SMT3 GG::CYC1t | This study | pMC_0486 |
| pCu415 pGAL1::SMT3 GG::CYC1t | This study | pFA_196 |
| pCu415 pGAL1::smt3 Q95P GG::CYC1t | This study | pFA_261 |
| pCu415 pGAL1::smt3 F37A GG::CYC1t | This study | pFA_264 |
| pCu415 pGAL1::smt3 F37A Q95P GG::CYC1t | This study | pFA_265 |
| *pCu415 pGAL1::smt3 F37A I39A GG::CYC1t* | This study | pFA_335 |
| *pCu415 pGAL1::smt3 F37A I39A Q95P GG::CYC1t* | This study | pFA_336 |
| *pCu426 pGAL1::smt3 Q95P GG::GFP::CYC1t* | This study | pFA_668 |
| *pCu426 pGAL1::smt3 F37A I39A Q95P GG::GFP::CYC1t* | This study | pFA_669 |
| *pGBKT7* | Clontech® | pMC_0385 |
| *pGBKT7 CSM1* | This study | pMC_0477 |
| *pGBKT7 LRS4* | This study | pMC_0479 |
| *pGBKT7 CDC14* | This study | pMC_0658 |
| *pGBKT7 SMT3 AA* | This study | pMC_0273 |
| *pGBKT7 smt3 F37A I39A AA* | This study | pMC_0275 |
| *pGADT7* | Clontech® | pMC_0386 |
| *pGADT7 HEH1 N (1-454aa)* | Capella et al., 2020 | pMC_0354 |
| *pGADT7 HEH1 C (730-834aa)* | Capella et al., 2020 | pMC_0355 |
| *pGADT7 NUR1 N (1-60aa)* | This study | pMC_0505 |
| *pGADT7 NUR1 M (167-240aa)* | This study | pMC_0507 |
| *pGADT7 NUR1 C (278-484aa)* | This study | pMC_0504 |
| *pGADT7 NUR1 278-332 (278-332aa)* | This study | pMC_0524 |
| *pGADT7 NUR1 278-430 (278-430aa)* | This study | pMC_0523 |
| *pGADT7 NUR1 333-484 (333-484aa)* | This study | pMC_0525 |
| *pGADT7 NUR1 431-484 (431-484aa)* | This study | pMC_0527 |
| *pGADT7 NUR1 333-430 (333-430aa)* | This study | pMC_0526 |
| *pGADT7 NUR1 S441D T446D (278-484aa)* | This study | pMC_0529 |
| *pGADT7 NUR1 S439D S441D T446D (278-484aa)* | This study | pMC_0535 |
| *pGADT7 NUR1 S441D T446D S449D (278-484aa)* | This study | pMC_0537 |
| *pGADT7 NUR1 S439D S441D T446D S449D (278-484aa)* | This study | pMC_0539 |
| *pGADT7 NUR1 S441A T446A (278-484aa)* | This study | pMC_0530 |
| *pGADT7 NUR1 S439A S441A T446A (278-484aa)* | This study | pMC_0536 |
| *pGADT7 NUR1 S441A T446A S449A (278-484aa)* | This study | pMC_0538 |
| *pGADT7 NUR1 S439A S441A T446A S449A (278-484aa)* | This study | pMC_0540 |
| *pGADT7 SMT3 GG* | This study | pMC_0276 |
| *pGADT7 SMT3 AA* | This study | pMC_0277 |
| *pGADT7 CDC48* | This study | pMC_0470 |
| *pGADT7 NPL4* | This study | pMC_0472 |
| *pGADT7 UFD1* | This study | pMC_0473 |
| *pGADT7 ufd1ΔSIM (1-355aa)* | This study | pMC_0474 |

**Supplementary Table S4:**
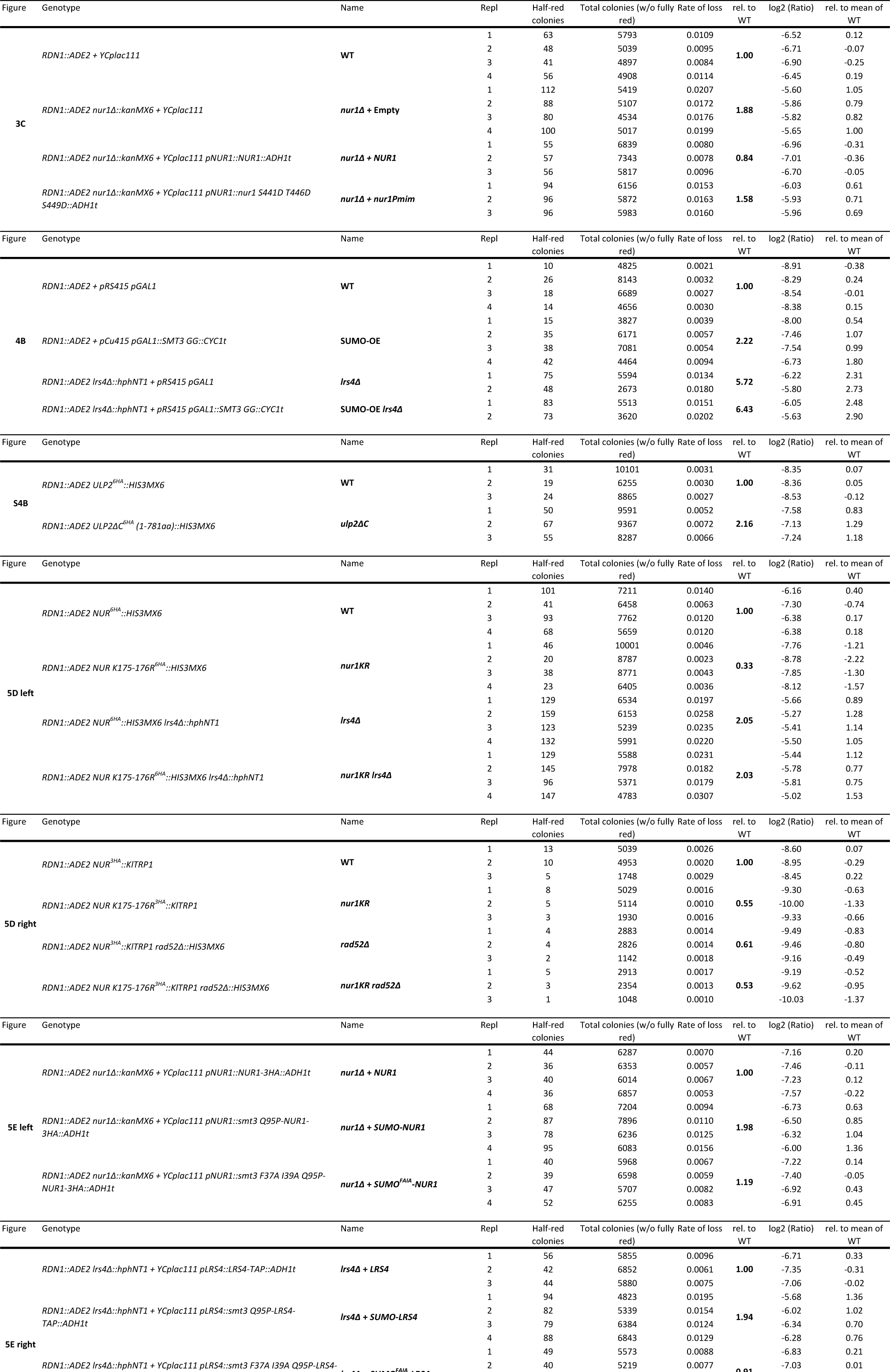
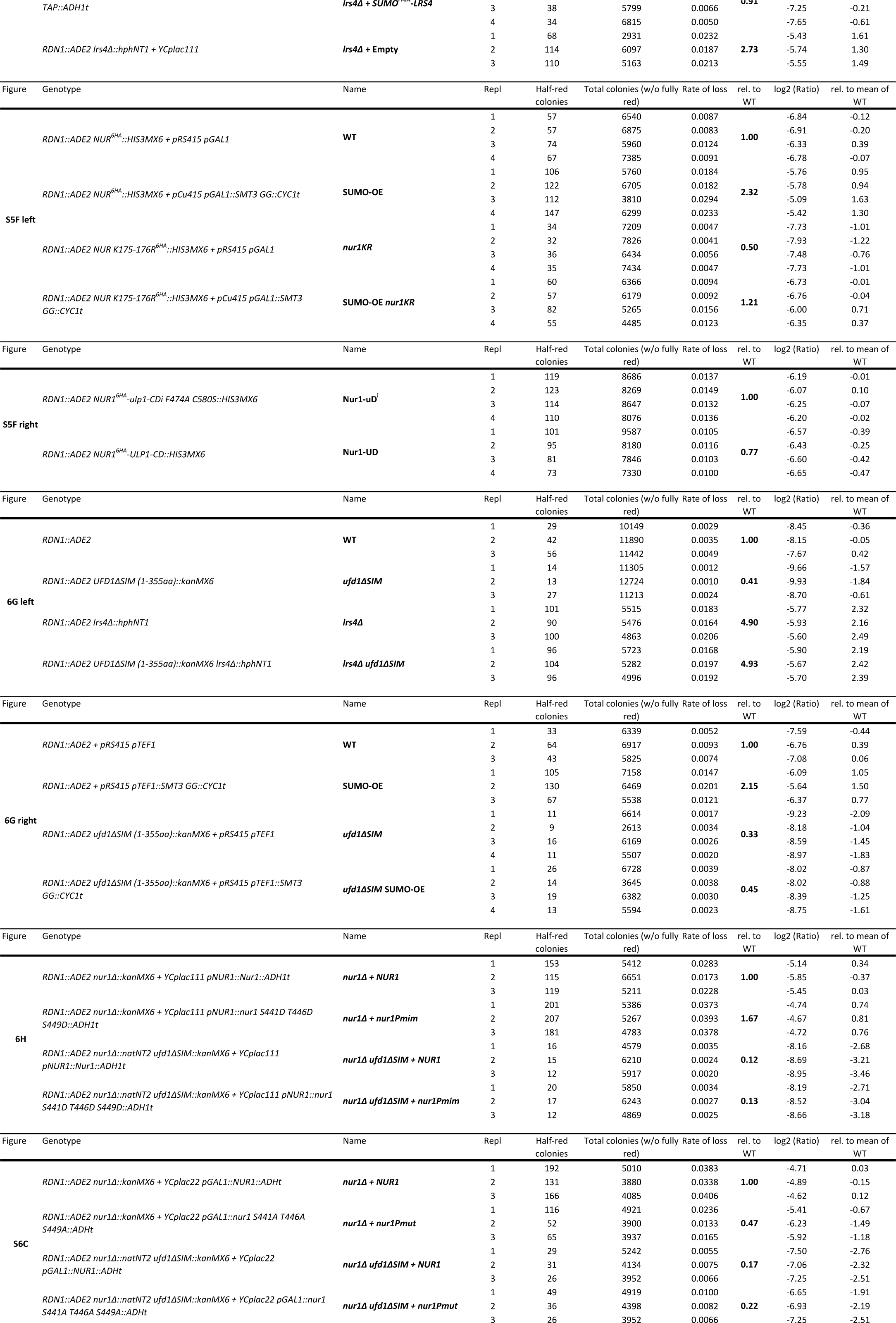
Rates of *ADE2* marker loss from rDNA repeats.

